# Intestinal Epithelial C/EBPβ Deficiency Impairs Colitis-Associated Tumorigenesis by Disrupting CXCL1/CXCL2/CXCL5-CXCR2-Mediated Neutrophil Infiltration

**DOI:** 10.1101/2025.06.25.661448

**Authors:** Mingyue Li, Xintong Wang, Wenjie Hu, Xiaohui Cheng, Qi Sun, Yongjie Wu, Zhen Huang, Jiangning Chen

## Abstract

Dysregulated transcription factors critically link chronic inflammation to oncogenesis in colitis-associated colorectal cancer (CAC), but their mechanistic roles remain incompletely understood. By integrating microarray and transcriptome sequencing data from ulcerative colitis (UC), colitis-associated cancer (CAC), and colorectal cancer (CRC) patients, we identify C/EBPβ as a key transcriptional regulator whose elevated expression inversely correlates with survival. In azoxymethane (AOM)/dextran sulfate sodium (DSS)-induced CAC models, intestinal epithelial C/EBPβ is upregulated during tumor progression, correlating with exacerbated tumor burden and neutrophil infiltration. Mice with intestinal epithelial-specific *Cebpb* deletion (*Cebpb*^ΔIEC^) show resistance to carcinogenesis, accompanied by reduced neutrophil infiltration and tumor growth. Mechanistically, C/EBPβ transcriptionally activates CXCR2 ligands (CXCL1, CXCL2, and CXCL5) to drive neutrophil recruitment. Pharmacological inhibition of CXCR2 phenocopies the anti-tumor effects of *Cebpb*^ΔIEC^ deletion, further validating this axis as a therapeutic target. Correlation analysis in patient tissues confirms positive relationships between C/EBPβ, CXCR2 ligands, and neutrophil infiltration, suggesting that targeting the C/EBPβ-CXCL1/2/5-CXCR2 axis may offer a novel strategy for CAC treatment.

## Introduction

Chronic inflammation is a well-established driver of carcinogenesis that significantly contributes to global disease burden and mortality [1, 2]. This relationship is particularly evident in colorectal cancer (CRC), the third-leading cause of cancer mortality worldwide [3], where inflammation-mediated oncogenesis plays a crucial role [4, 5]. The connection is most pronounced in colitis-associated colorectal cancer (CAC), which develops in 2.4% of inflammatory bowel disease (IBD) patients within 20 years of diagnosis [6]. Notably, while only 5.2% (95% CI 4.3-6.1%) of CRC cases originate from overt inflammation [7, 8]. these patients exhibit markedly worse 5-year survival rates compared to sporadic CRC (18.3% vs. 64.1%, P<0.001)[9], primarily due to therapeutic resistance [10]. The molecular pathogenesis of CAC involves multiple mechanisms: (i) early activation of oncogenic pathways, evidenced by the 3.2-fold higher frequency of KRAS^G12D^ mutations compared to sporadic CRC [11] ; (ii) dysregulation of key tumor suppressors including TP53 and DCC [12–14]; and (iii) establishment of a persistent inflammatory microenvironment. Despite these advances in understanding CAC development, the precise mechanistic links between chronic inflammation and CAC progression remain incompletely characterized, particularly regarding the transcriptional regulation of this process.

CAC pathogenesis involves transcriptional reprogramming linking inflammation to malignancy. Central to this process are transcription factors that coordinately regulate gene networks governing immune homeostasis and cellular proliferation. Key players include early pathogenic mediators such as *TP53* mutations and constitutive TNF/NF-κB activation [15, 16], along with dysregulated cofactors like NFKBIZ [17, 18] and epithelial-specific defects including SMAD4 loss (elevating CCL20-CCR6 axis activity) [19] and ITF2 deficiency (promoting p65 nuclear translocation) [17].

Among transcriptional regulators linking inflammation to carcinogenesis, CCAAT/enhancer-binding protein β (C/EBPβ) has emerged as a critical mediator. This conserved transcription factor contains a basic leucine zipper (bZIP) domain enabling DNA binding through homo- or heterodimerization [20], and gives rise to three functionally distinct isoforms: liver enriched activator protein (LAP1/LAP2) and liver enriched inhibitory protein (LIP) [21]. The activity of C/EBPβ is finely tuned through post-translational modifications that regulate its subcellular localization and protein interactions. Physiologically, C/EBPβ orchestrates diverse programs ranging from cell proliferation and differentiation to immune responses [22], while pathologically it integrates key inflammatory signals including MAPK/NF-κB and JAK/STAT3 pathways [23]. Our previous study demonstrated its oncogenic role through C/EBPβ-miR-223-RASA1 axis in colorectal cancer [24]. Notably, despite these advances in understanding C/EBPβ’s general functions, its epithelium-specific mechanisms in CAC pathogenesis - particularly regarding neutrophil recruitment through chemokine regulation - remain poorly characterized.

In this study, our multi-omics analysis of ulcerative colitis (UC), CRC, and CAC patient datasets identified C/EBPβ as a key transcription factor in CAC pathogenesis. Clinical samples and azoxymethane (AOM)/dextran sulfate sodium (DSS)-induced mouse models confirmed intestinal epithelial C/EBPβ drives CAC progression, with genetic deletion reducing inflammation and tumor burden. Mechanistically, C/EBPβ promotes neutrophil recruitment through transcriptional activation of CXCR2 ligands (CXCL1/2/5), thereby establishing a pro-tumorigenic inflammatory microenvironment. CXCR2 inhibition attenuated neutrophil infiltration, and human tissue analyses confirmed C/EBPβ-CXCR2 ligand correlations. Our study identifies epithelial C/EBPβ as a previously unrecognized mechanistic link connecting inflammation to CAC progression, specifically via transcriptional activation of CXCR2 ligands. The C/EBPβ-CXCL1/2/5-CXCR2 axis represents a novel therapeutic target for preventing inflammation-driven colorectal carcinogenesis.

## Materials and Methods

### Clinical samples

Clinical samples of UC, CAC and CRC from public datasets (GEO: GSE87466, GSE37283; TCGA-COAD/READ) were analyzed (Supplementary Tables S1-3). Human tissue samples were analyzed from: (1) 180 CRC patient-matched pairs (Outdo Biotech; Supplementary Table S4), (2) 5 UC and 5 CAC cases with matched controls, plus 10 normal biopsies (Nanjing Jinling Hospital; Supplementary Table S5), and (3) 64 additional CRC cases (Nanjing Jinling Hospital; Supplementary Table S6). The study was conducted with ethics approval (Jinling Hospital No. 2021DZSKT-YBB-008) and informed consent.

### Reagents

Reagents included AOM and SB225002 (Sigma, St. Louis, USA), DSS (MP Biomedicals, Solon, USA), *CEBPB* siRNA from Thermo Fisher (Grand Island, USA) (Supplementary Table S7), and a pcDNA3.1-*CEBPB* plasmid (Real Gene, Nanjing, China). Luciferase reporters (pGL3 vectors, Promega, Madison, USA) contained 2 kb promoters of CXCL1/2/5 with wild-type or mutated *CEBPB* sites. For CXCL1, the sequence CTTTCAAAAT was altered to ACGGTCCCCA; for CXCL2, GTTTCACAAC was changed to AGGGTCTCCT; and for CXCL5, TCTTGCTCCAT was replaced with TCAATTTTTAT. Biotin-labeled and unlabeled probes were synthesized by GeneBio Co., Ltd (Shanghai, China; Supplementary Table S7). Recombinant human C/EBPβ protein was purchased from Proteintech (Wuhan, China).

### Animal studies

Male C57BL/6J mice (6-8 weeks old) were purchased from GemPharmatech (Nanjing, China). Intestinal epithelial-specific *Cebpb* knockout (*Cebpb*^ΔIEC^) mice were generated by crossing B6.129S1-*Cebpb*^tm1Rcsm^/Mmnc mice with B6.Cg-Tg(Vil1-cre)1000Gum/J strains. Mice were housed under SPF conditions. All animal procedures were approved by Nanjing University Animal Ethics Committee (IACUC-2006015).

For CAC induction, 8-week-old mice received 10 mg/kg AOM intraperitoneally, followed by three cycles of 2.5% DSS (1 week) and normal water (2 weeks). SB225002 (10 mg/kg, twice weekly) was used to assess neutrophil infiltration. Mice were randomized into five groups (n = 5/group): 1. wild-type (WT) controls receiving intraperitoneal saline and normal drinking water; 2. *Cebpb*^ΔIEC^ mice treated with AOM/DSS; 3. *Cebpb*^ΔIEC^ mice treated with AOM/DSS/SB225002; 4. WT mice treated with AOM/DSS; 5. WT mice treated with AOM/DSS/SB225002. On day 63, colons were collected for histopathology, qRT-PCR, Western blot, flow cytometry, RNA-seq, and ELISA.

### Histopathological analysis

Human and mouse tissues were fixed, paraffin-embedded and sectioned. H&E staining was performed according to manufacturer protocols (Leagene Biotechnology), with mouse tumors classified as low-grade dysplasia, high-grade dysplasia, or invasive carcinoma using established criteria [25]. For immunofluorescence, antigen-retrieved sections were incubated overnight at 4°C with Alexa Fluor-conjugated antibodies (Ly-6G, CD11b) followed by nuclear counterstaining with DAPI (Beyotime), with appropriate isotype controls. Images were acquired using a confocal microscope (ZEISS LSM 980). Immunohistochemical staining involved overnight incubation at 4°C with primary antibodies (C/EBPβ, CXCL1/2/5 and CD66b) after antigen retrieval and peroxidase blocking, followed by 45-minute room temperature incubation with HRP-conjugated secondary antibodies, DAB development, and hematoxylin counterstaining, with imaging performed on a Nikon Eclipse Ni-E upright microscope. Secondary antibodies alone (IHC) were included to assess nonspecific binding.

Quantitative analysis included: human C/EBPβ immunoreactive scoring (IRS 0-12 combining intensity and positivity) and percentage DAB-positive area for mouse/human markers, and neutrophil cell density (human: CD66b^+^ within total cells; mouse: CD11b^+^ Ly-6G^+^ within CD11b^+^ myeloid cells). Two blinded pathologists performed all analyses using ImageJ (≥5 fields/sample). Antibody’s information is detailed in Supplementary Table S8.

### Cell culture and functional assays

Human colorectal adenocarcinoma Caco-2 and promyelocytic leukemia HL-60 cells (STR-profiled, mycoplasma-free) were cultured following protocols. HL-60 cells were differentiated with 5 μM retinoic acid (3 days). Transfections with pcDNA3.1-*CEBPB*/*CEBPB* siRNA were assessed by qRT-PCR (24 h) and ELISA (CXCL1/2/5; 48 h). Luciferase assays used co-transfected reporter plasmids and β-galactosidase controls (Promega) to investigate the binding of *CEBPB* to the promoters of CXCL1/2/5. For chemotaxis, neutrophil-like HL-60 cells (SB225002-treated) were exposed to Caco-2 supernatants in Transwells (3 μm pores; CORNING, Corning, USA), with migrated cells quantified after crystal violet staining.

### Flow cytometry analysis

Primary colonic immune cells were isolated through sequential enzymatic digestion using EDTA/DTT (Sigma), followed by collagenase D/DNase I/hyaluronidase (Sigma) treatment and Percoll (Sigma) gradient centrifugation [26]. After isolation, cells were blocked with Fc-specific antibodies, stained with 7-AAD viability marker prior to analysis (Attune NxT, FlowJo v10). Antibody details are provided in Supplementary Table S8.

### Molecular biology assays

RNA analysis: Total RNA was extracted with TRIzol (Thermo Fisher), reverse-transcribed (TaKaRa kit, Shiga, Japan), and analyzed by qRT-PCR (StepOnePlus) with β-actin as the endogenous control (primers listed in Supplementary Table S9).

Protein analysis: During colonic immune cell isolation, epithelial and stromal fractions were separately collected. Protein lysates were extracted using RIPA buffer (Beyotime, Shanghai, China), quantified with a BCA assay (Vazyme, Nanjing, China), and analyzed by immunoblotting. Fraction purity was confirmed through marker absence: epithelial fractions showed no detectable CD45 (leukocyte marker) or α-SMA (stromal marker), while stromal fractions were negative for Pan Cytokeratin (epithelial marker).

Chromatin immunoprecipitation (ChIP): DNA-protein complexes were immunoprecipitated using anti-C/EBPβ antibody (ChIP kit, Millipore, Billerica, USA), followed by qPCR analysis of precipitated DNA.

Electrophoretic Mobility Shift Assay (EMSA): The LightShift Chemiluminescent EMSA Kit (Thermo Fisher Scientific) was used for EMSA analysis. Biotin-labeled DNA probes containing putative C/EBPβ binding sites were incubated with recombinant human C/EBPβ protein to form DNA-protein complexes. Reaction mixtures were resolved on 6% non-denaturing polyacrylamide gels in 0.5× TBE buffer (100 V, 60 min), then transferred to nylon membranes (380 mA, 30 min) and UV-crosslinked (120 mJ/cm², 1 min) prior to chemiluminescent detection.

Chemokine quantification: Tissue supernatant levels of CXCL1 (EK296, Multi Sciences, Hangzhou, China), CXCL2 (EK2142, Multi Sciences), and CXCL5 (EK0919, Boster) were measured by ELISA according to manufacturers’ protocols.

### RNA sequencing analysis

Total RNA was extracted from the mid-distal colon tissues using TRIzol reagent (Thermo Fisher Scientific), following the manufacturer’s guidelines. To remove any genomic DNA contamination, the RNA was treated with RNase-free DNase. cDNA library was prepared and sequenced on an Illumina NovaSeq 6000 platform (San Diego, USA) by CapitalBio Corporation (Beijing, China). Differential expression analysis was performed using DESeq2 (v1.1), with genes showing |log2(Fold Change (FC)| > 1 and adjusted P-value < 0.05 considered statistically significant. The raw sequencing data have been deposited in the NCBI Gene Expression Omnibus under accession number GSE264497.

### Bioinformatics analysis

We performed integrated analysis of microarray datasets from GEO (GSE87466: 21 normal, 87 UC; GSE37283: 5 normal, 11 CAC) and TCGA-COAD/READ (51 normal, 383 CRC) to identify consistently dysregulated transcription factors in UC, CAC and CRC. All samples were processed using Affymetrix HT HG-U133+ PM Arrays. Data were normalized using Robust Multi-array Average (RMA) and batch-corrected with the ComBat algorithm. Candidate transcription factors were screened using Animal Transcription Factor Database (Animal TFDB v3.0), with differential expression defined as |log2FC| > 1 and P < 0.05 (DEseq2). Results were visualized using ggplot2 and pheatmap R packages. Additionally, immune cell infiltration in CRC and CAC tissues was assessed via CIBERSORTx website (http://cibersortx.standford.edu/) [27, 28]. For functional annotation, we conducted KEGG pathway analysis (DAVID) (https://david.ncifcrf.gov/) [29] and GSEA using the Broad Institute platform [30]. Statistical significance thresholds were set at P < 0.05 for KEGG and FDR < 0.25 for GSEA.

### Statistical analysis

Data are presented as mean ± standard deviation (SD). All analyses were performed using GraphPad Prism 9.0 (GraphPad Software Inc., La Jolla, CA, USA). Normality was assessed using Shapiro-Wilk tests, while homogeneity of variance was evaluated with Brown-Forsythe tests. Parametric tests were applied to normally distributed data with equal variances: unpaired two-tailed t-tests for two-group comparisons and one-way ANOVA with Bonferroni correction for multiple comparisons. Non-normally distributed data were analyzed using Mann-Whitney U tests (two groups) or Kruskal-Wallis tests with Dunn’s post hoc correction (multiple groups). For normally distributed data with unequal variances, Welch’s t-test was employed. Linear relationships between biomarkers (C/EBPβ, CXCL1/2/5, CD66b) were quantified using Pearson correlation coefficients. Statistical significance was defined as *P* < 0.05 (two-tailed), with exact *P*-values reported except when *P* < 0.001.

## Results

### Identification of dysregulated transcription factors in UC, CAC, and CRC pathogenesis

Through integrated analysis of GEO and TCGA datasets (Supplementary Tables S1-3), we identified five transcription factors with consistent dysregulation patterns across ulcerative colitis (UC), colitis-associated cancer (CAC), and colorectal cancer (CRC). Venn analysis comparing UC versus healthy controls, CAC versus controls, and CRC versus normal adjacent tissue (NAT) revealed *CEBPB* and *FOXQ1* as significantly upregulated (log2FC > 1, P < 0.05), while *ISX*, *NR3C2*, and *SATB2* were downregulated (log2FC < -1, P < 0.05) in all three conditions (Figure 1A and Supplementary Data 1). Heatmap visualization and histograms confirmed these distinct expression profiles (Figure 1B-D and Supplemetary Figure S1A-L). *CEBPB* exhibited particularly striking patterns, showing progressive overexpression from UC to CAC to CRC (Figure 1E-G). Clinical correlation analysis demonstrated that elevated *CEBPB* expression significantly predicted poorer survival in CRC patients (P < 0.01, Figure 1H), whereas the other four transcription factors showed no prognostic significance (Supplementary Figure S1M-P). These findings establish CEBPB as a consistently upregulated transcription factor throughout the inflammation-cancer transformation cascade and a potential prognostic marker in CRC.

**Figure 1.**
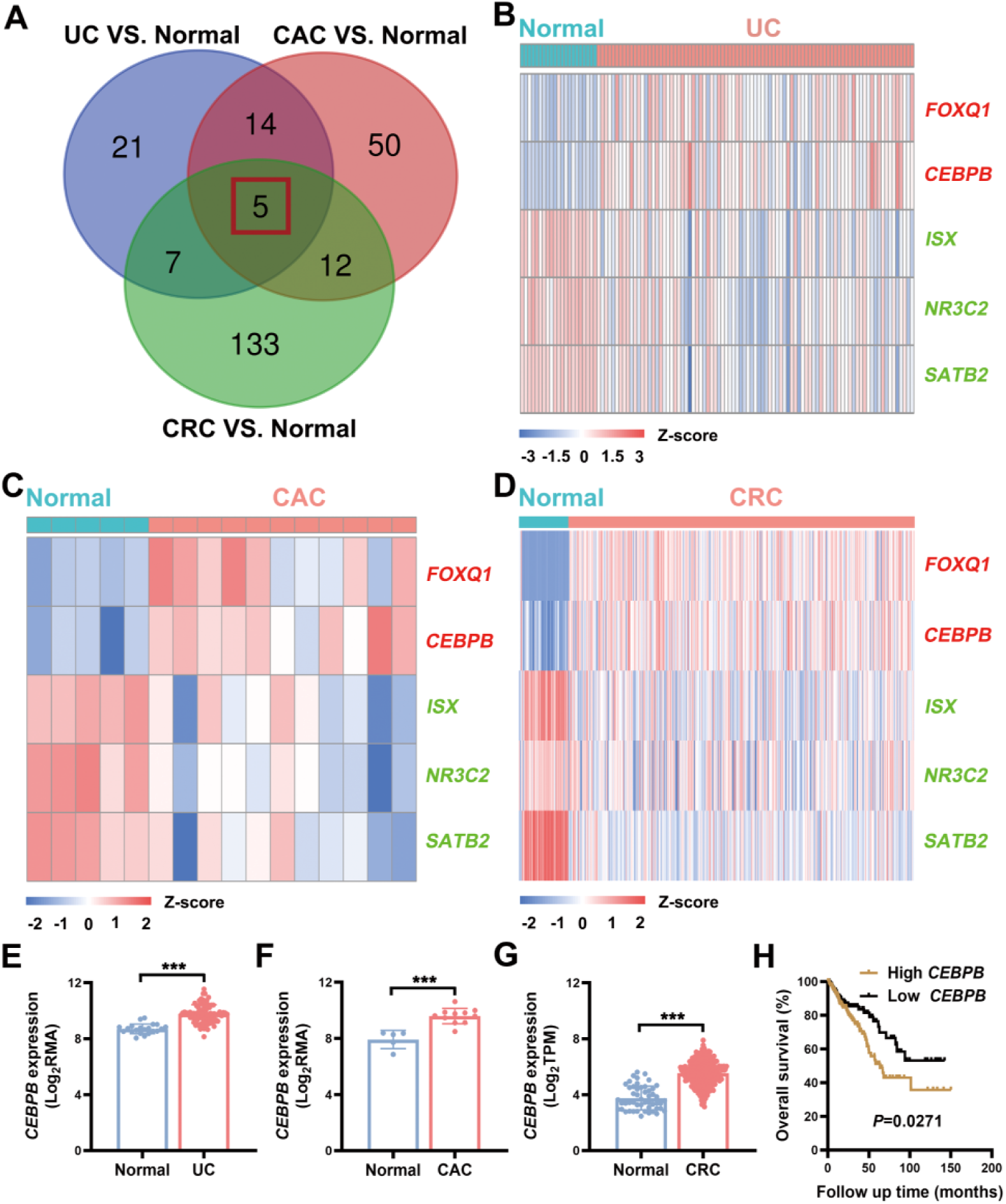
Transcriptional landscape of transcription factors in patients with ulcerative colitis (UC), colitis-associated cancer (CAC), and colorectal cancer (CRC). (A) Venn diagram of differentially expressed transcription factors from GEO/TCGA databases. (B-D) Heatmaps of 5 consistently dysregulated transcription factors (red: upregulated; green: downregulated). (E-G) *CEBPB* mRNA expression in different patient cohorts: UC patients (GSE87466: 21 healthy individuals vs 87 UC; Welch’s two-tailed t-test); CAC patients (GSE37283: 5 healthy individuals vs 11 CAC; unpaired t-test); and CRC patients (TCGA: 51 normal adjacent tumor (NAT) vs 383 tumors; Mann-Whitney U test). (H) Kaplan-Meier survival analysis of 376 CRC patients stratified by *CEBPB* mRNA expression (TCGA). Data are expressed as mean ± SD. \*\*\**P* < 0.001.

### C/EBP**β** overexpression correlates with disease progression and poor prognosis in CAC and CRC

Immunohistochemistry analysis of CAC and UC patient tissues demonstrated predominant C/EBPβ expression in colonic epithelial cells (Figure 2A, B, Supplementary Figure S2A, B and Supplementary Table S5). Quantitative assessment revealed progressively increasing C/EBPβ levels from normal mucosa to adjacent non-cancerous tissues (CAC-AT) and finally to CAC lesions (P<0.01). This escalation was confirmed at the transcriptional level by qRT-PCR (Figure 2C). In sporadic CRC, tissue microarray (TMA) analysis of 180 cases showed similar C/EBPβ upregulation (Figure 2D, E and Supplementary Table S4). Importantly, we observed a significant positive correlation between C/EBPβ expression levels and TNM staging (Figure 2E), with elevated C/EBPβ predicting reduced overall survival (Figure 2F). Western blot and qRT-PCR validation in independent CRC cohorts (Supplementary Table S6) confirmed consistent C/EBPβ overexpression (Figure 2G-I), corroborating our bioinformatic survival analysis (Figure 1H). These results establish C/EBPβ as a clinically significant biomarker showing stage-dependent overexpression and prognostic value in both CAC and sporadic CRC pathogenesis.

**Figure 2.**
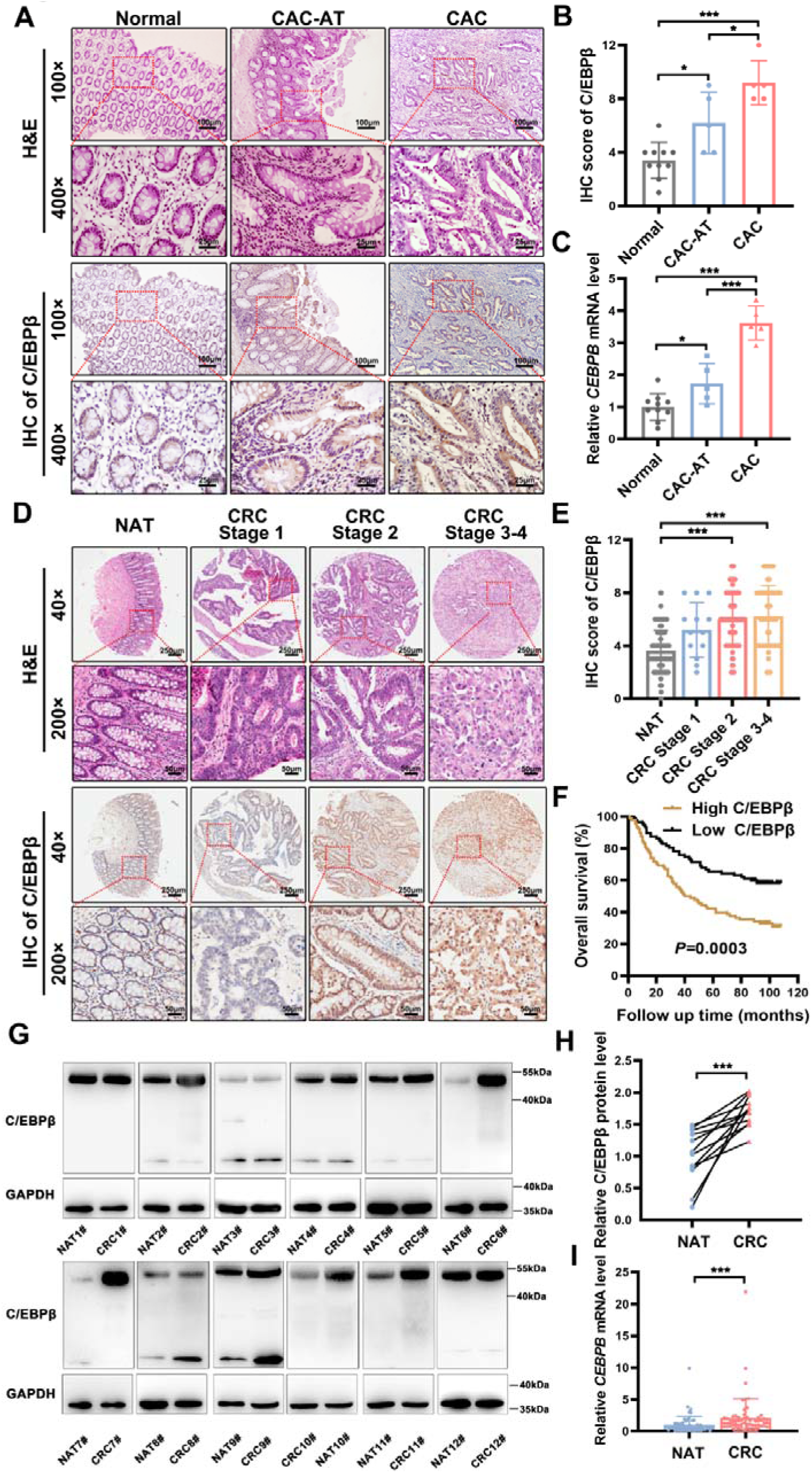
Clinical significance of elevated C/EBPβ in colorectal cancer progression. (A) Representative hematoxylin and eosin (H&E) and immunohistochemistry (IHC) staining of C/EBPβ in normal colon, CAC-adjacent (CAC-AT), and CAC tissues (100×: 100 μm scale; 400×: 25 μm scale). (B-C) Quantitative analysis of (B) C/EBPβ protein (IHC) and (C) C*EBPB* mRNA levels (normal: n = 10; CAC: n = 5; one-way ANOVA with Bonferroni correction). (D) CRC tissue microarrays (n = 180) showing C/EBPβ expression (40×: 250 μm scale; 200×: 50 μm scale). (E) C/EBPβ IHC scores across CRC Stages (Kruskal-Wallis with Dunn’s test). (F) Kaplan-Meier survival analysis of 180 CRC patients by C/EBPβ expression. (G-H) Western blot analyses of C/EBPβ expression in randomly selected CRC patients (n = 12). Protein levels were quantified by densitometry (normalized to GAPDH) using ImageJ software. Statistical analysis was conducted using the Mann-Whitney U test. (I) qRT-PCR was employed to evaluate *CEBPB* mRNA levels in randomly selected CRC patients (n = 64). Statistical analysis was performed using the Mann-Whitney U test. Data are expressed as mean ± SD. \**P* < 0.05, \*\*\**P* < 0.001.

### Intestinal epithelial *Cebpb* deletion mitigates tumorigenesis in AOM/DSS-induced murine CAC models

We successfully established the AOM/DSS-induced CAC model in mice. Representative colon images demonstrate the morphological differences between untreated WT mice and AOM/DSS-treated WT mice (Figure 3A). Quantitative analysis revealed significantly elevated *Cebpb* mRNA levels in colonic lesions of AOM/DSS-treated mice compared to controls (Figure 3B). Western blot analysis confirmed this upregulation was specifically localized to intestinal epithelial cells (Figure 3C), suggesting their central role in C/EBPβ-mediated pathogenesis. To functionally characterize C/EBPβ’s contribution, we generated intestinal epithelial-specific *Cebpb*-knockout mice (*Cebpb*^Δ*IEC*^) through Villin-cre mediated recombination of floxed alleles (Supplementary Figure S3). The *Cebpb*^ΔIEC^ mice showed remarkable resistance to AOM/DSS-induced tumorigenesis with a 62.5% reduction in tumor number compared to AOM/DSS-induced ittermate controls (Figure 3D, E). Histopathological examination demonstrated that *Cebpb*^ΔIEC^ mice showed near-normal intestinal architecture with significantly attenuated inflammatory cell infiltration and developed low-grade dysplasia following AOM/DSS treatment (Figure 3F and Supplementary Table S10). The epithelial-specific deletion of C/EBPβ was verified through immunohistochemical staining (Figure 3F, G), transcriptional analysis by qRT-PCR (Figure 3H), and protein level assessment by Western blot (Figure 3I). These findings definitively establish intestinal epithelial C/EBPβ as a critical molecular driver of colitis-associated carcinogenesis.

**Figure 3.**
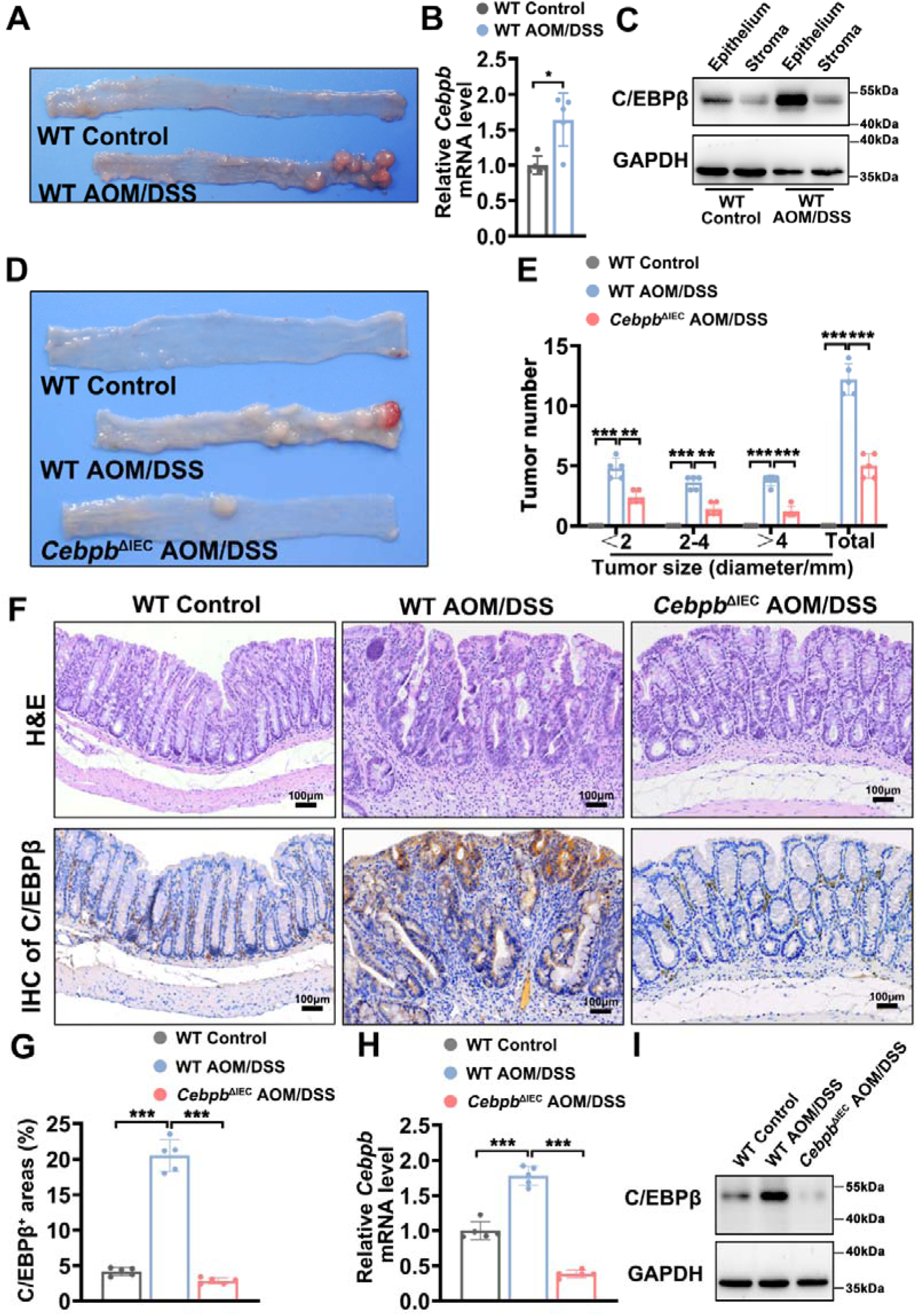
Effects of C/EBPβ deletion in intestinal epithelial cells on tumorigenesis in AOM/DSS-induced murine models. (A) Representative images of colons from untreated wild-type (WT) control mice and AOM/DSS-treated WT mice. (B) *Cebpb* mRNA levels in colonic tissue from WT control and AOM/DSS-treated WT mice were quantified by qRT-PCR. Statistical analysis was performed by Mann-Whitney U test; n = 5 mice/group. (C) Western blot analysis of C/EBPβ protein expression in colonic epithelium and stroma of WT control and AOM/DSS-treated WT mice. Protein levels were normalized to GAPDH. (D) Representative images of colons from three experimental groups: (i) WT control mice, (ii) AOM/DSS-treated WT mice (WT AOM/DSS), and (iii) AOM/DSS-treated mice with intestinal epithelial cell-specific *Cebpb* knockout (*Cebpb*^ΔIEC^ AOM/DSS). Tumor number and size were assessed; n = 5 mice/group. (E) Colon tumors were quantified by size stratification (>2 mm, measured by calipers; <2 mm, assessed by dissection microscope). Data were analyzed by one-way ANOVA with Bonferroni’s multiple comparisons test; n = 5 mice/group. (F) Representative H&E and IHC staining of C/EBPβ in colon tissues from WT control, WT AOM/DSS and *Cebpb*^ΔIEC^ AOM/DSS mice. Scale bar = 100 μm. (G) Quantification of C/EBPβ-positive staining areas in colon tissues from WT control, WT AOM/DSS and *Cebpb*^ΔIEC^ AOM/DSS mice by ImageJ software. (H-I) *C/EBP*β mRNA levels (qRT-PCR) and protein expression (Western blot) in colon tissues from WT control, WT AOM/DSS and *Cebpb*^ΔIEC^ AOM/DSS mice. Statistical analysis was performed by Kruskal-Wallis test with Dunn’s multiple comparisons test. n = 5 mice/group for qRT-PCR. Data are presented as mean ± SD. ns: not significant, \**P* < 0.05, \*\**P* < 0.01, \*\*\**P* < 0.001.

### Intestinal epithelial C/EBP**β** promotes neutrophil recruitment during malignant transformation

Flow cytometric analysis of AOM/DSS-treated mice revealed substantial infiltration of innate immune cells, particularly neutrophils and macrophages, in colonic tissues. Targeted deletion of *Cebpb* specifically in intestinal epithelial cells (*Cebpb*^ΔIEC^) resulted in a marked 78% reduction in neutrophil recruitment compared to AOM/DSS-treated wild-type mice (Figure 4A-C and Supplementary Figure S4). This selective effect on neutrophil infiltration was further supported by clinical data analysis using the CIBERSORT algorithm, which demonstrated significantly elevated neutrophil signatures in transcriptomic profiles from patients with ulcerative colitis (UC), colitis-associated cancer (CAC), and colorectal cancer (CRC) (Figure 4D-I). The concordance between murine experimental data and human clinical observations strongly implicates intestinal epithelial C/EBPβ as a specific regulator of neutrophil recruitment during the inflammation-to-cancer transition.

**Figure 4.**
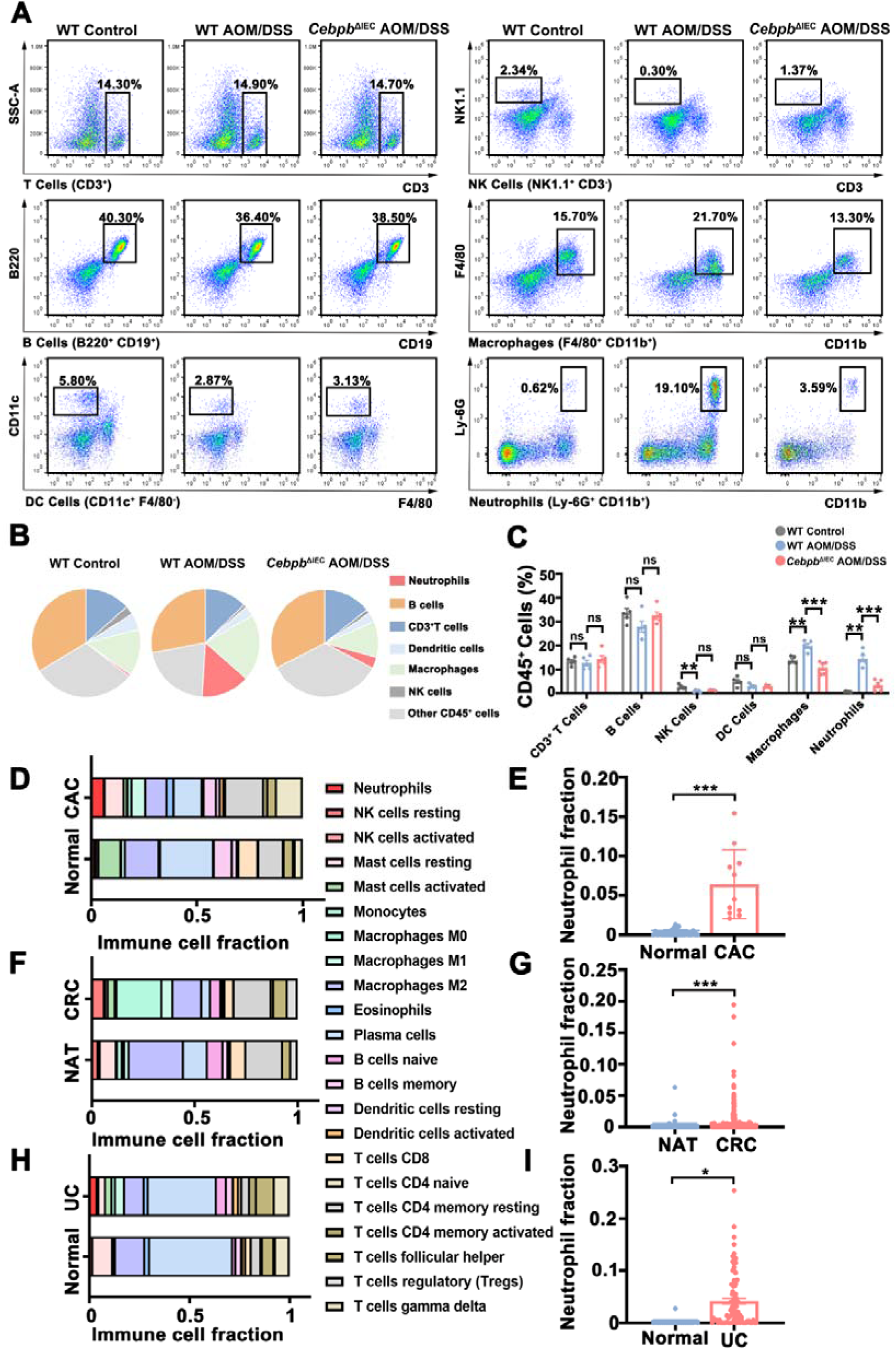
*Cebpb* deletion in intestinal epithelial cells reduces neutrophil infiltration in AOM/DSS-treated mice. (A-C) Immune cell subsets in colon tissues from WT control, WT AOM/DSS, and *Cebpb*^ΔIEC^ AOM/DSS mice were quantified by flow cytometry (n = 3 mice/group). Data analyzed by one-way ANOVA with Bonferroni’s correction. (D-I) Bioinformatics analysis of neutrophil-associated genes in human datasets (Mann-Whitney U test): For panels D-E, data were obtained from 26 healthy individuals and 11 CAC patients (GSE87466, GSE37283). For panels F-G, data were obtained from 41 normal adjacent tumor tissues (NAT) and 216 CRC samples (TCGA-COAD/READ). For panels H-I, data were collected from 21 healthy individuals and 87 UC patients (GSE87466 dataset). Data are expressed as mean ± SD. ns: not significant, \**P* < 0.05, \*\**P* < 0.01, \*\*\**P* < 0.001.

### C/EBP**β** transcriptionally activates CXCR2 ligands to mediate neutrophil infiltration in CAC pathogenesis

Transcriptomic profiling of colonic tissues from WT, AOM/DSS-treated, and *Cebpb*^ΔIEC^ AOM/DSS mice revealed significant enrichment of cytokine-cytokine receptor interaction pathways (Figure 5A, B and Supplementary Data 2-3). Gene Set Enrichment Analysis (GSEA) confirmed strong positive correlation of this pathway in WT AOM/DSS mice versus controls, while *Cebpb* deletion reversed this trend (Figure 5C, D). Among 36 differentially expressed genes identified by intersection analysis (Supplementary Figure S5 and Supplementary Table S11), CXCR2 ligands (*Cxcl1*, *Cxcl2*, and *Cxcl5*) and *Cxcr2* itself were markedly upregulated in WT AOM/DSS mice but significantly attenuated in C*ebpb*^ΔIEC^ mice (Figure 5E, F). This regulation was validated at both transcriptional (qRT-PCR) and protein (ELISA) levels (Figure 5G, H). The leading edge subset of genes is detailed in Supplementary Data 4.

**Figure 5.**
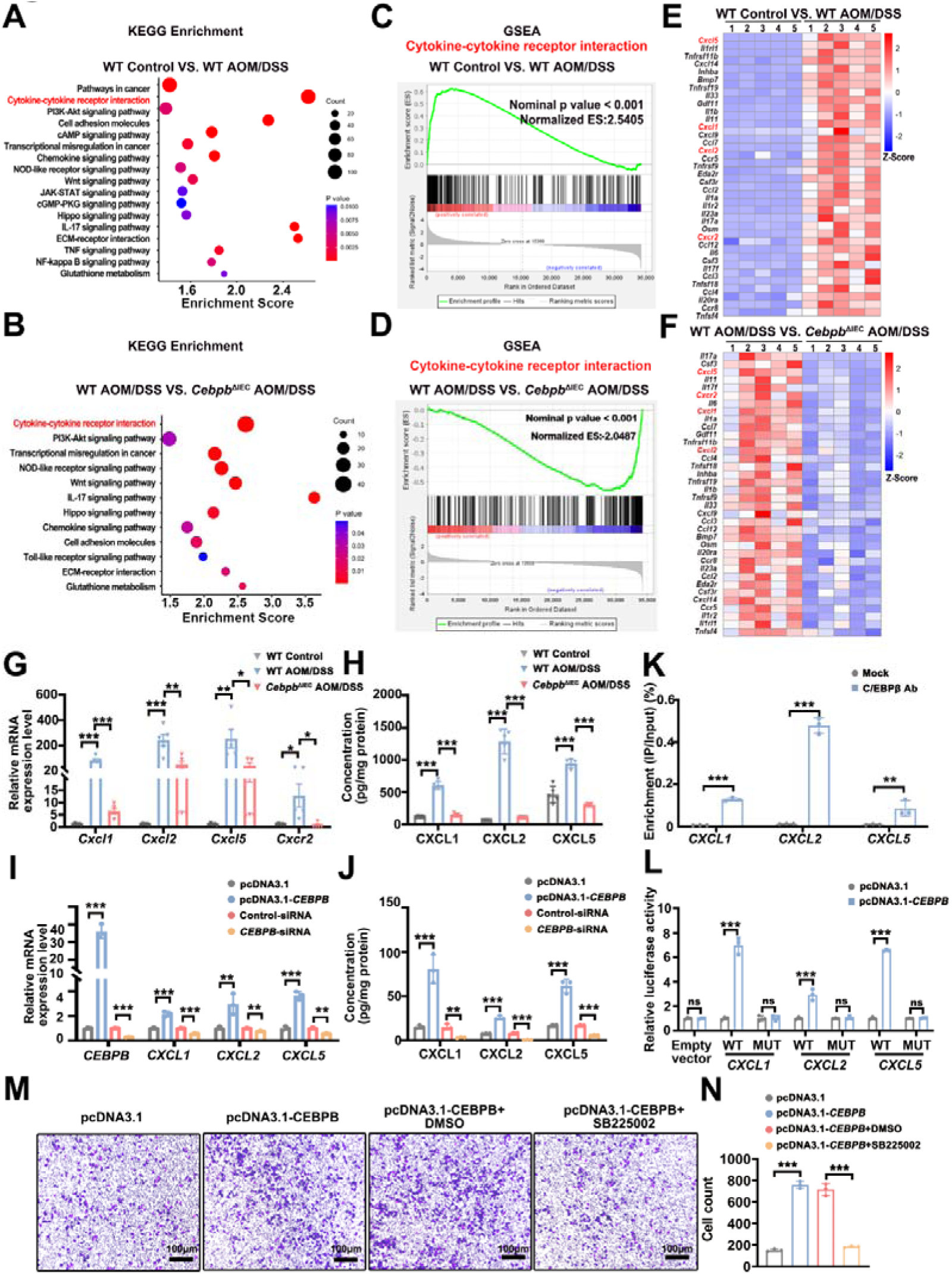
C/EBPβ mediates neutrophil recruitment through transcriptional upregulation of CXCR2 ligands. (A-D) KEGG and gene set enrichment analysis (GSEA) of differentially expressed genes (DEGs) in colon tissues from WT control, WT AOM/DSS, and *Cebpb*^ΔIEC^ AOM/DSS mice (n = 5 mice /group). (E-F) Heatmap showing 36 significantly enriched DEGs from the cytokine-cytokine receptor interaction pathway (Z-scores shown by color gradient). (G-H) CXCR2 ligand (CXCL1/2/5) and receptor levels in colon tissues measured by qRT-PCR (G; n = 5) and ELISA (H; n = 5. Statistical analysis was performed by one-way ANOVA with Bonferroni’s correction. (I-J) CXCL1/2/5 expression in Caco-2 cells with *CEBPB* overexpression (pcDNA3.1-*CEBPB*) or knockdown (*CEBPB*-siRNA), compared to their respective controls (one-way ANOVA with Bonferroni’s correction). (K) ChIP-qPCR confirming C/EBPβ binding to CXCL1/2/5 promoters (anti-C/EBPβ antibody vs. Input/IgG controls; unpaired t-test). (L) Luciferase reporter assays of *CEBPB-*responsive promoter activity (wild-type vs. mutant CXCL1/2/5 promoters) in *CEBPB*-overexpressing Caco-2 cells (unpaired t-test). (M-N) Representative images showing neutrophil migration toward supernatants from *CEBPB*-modulated Caco-2 cells (SB225002-pretreated; 100× magnification, scale bar = 100 μm; one-way ANOVA with Bonferroni’s correction) n = 3 biological replicates for panels I-N. Data are expressed as mean ± SD. ns: not significant, \**P* < 0.05, \*\**P* < 0.01, \*\*\**P* < 0.001.

Mechanistic studies in Caco-2 cells demonstrated that C/EBPβ directly activates CXCL1/2/5 transcription. Overexpression of C/EBPβ increased CXCR2 ligand expression, while knockdown reduced it (Figure 5I-J). Our integrated computational (JASPAR) and experimental (ChIP-qPCR) analyses identified C/EBPβ binding sites in the promoter regions of CXCL1 (chr4: 73868520-73868724), CXCL2 (chr4 complement: 74100928-74100673), and CXCL5 (chr4 complement: 74000013-73999933) (Figure 5K). Functional validation through luciferase reporter assays confirmed that mutation of these sites abolished C/EBPβ-mediated transactivation (Figure 5L), while EMSA demonstrated direct C/EBPβ binding *in vitro* (Supplementary Figure S6). Using CXCR2-deficient Caco-2 cells [31], we further showed that supernatants from C/EBPβ-overexpressing cultures induced robust neutrophil chemotaxis - an effect potently inhibited by the selective CXCR2 antagonist SB225002 (Figure 5M, N) [32]. These results establish a C/EBPβ→CXCL1/2/5→CXCR2 signaling axis whereby intestinal epithelial C/EBPβ transcriptionally drives neutrophil recruitment during colitis-associated carcinogenesis.

### Pharmacological blockade of C/EBP**β**-CXCR2 axis attenuates neutrophil-driven tumorigenesis in CAC

Building on previous reports demonstrating that neutrophil depletion reduces AOM/DSS-induced tumor burden [32], we investigated whether targeting the C/EBPβ-CXCR2 axis could similarly impair carcinogenesis. The CXCR2 antagonist SB225002 significantly reduced tumor burden in AOM/DSS-treated mice, as evidenced by decreased tumor number, size and histologic grade (Figure 6A, B and Supplementary Table S12), recapitulating the protective effects observed in *Cebpb*^ΔIEC^ mice. Flow cytometric and immunofluorescence analysis confirmed that SB225002 treatment reduced colonic neutrophil infiltration, comparable to genetic *Cebpb* ablation (Figure 6C-G). Notably, combined SB225002 treatment and *Cebpb* deletion provided no additional benefit, indicating both interventions target the same pathway. These findings demonstrate that intestinal epithelial C/EBPβ promotes CAC progression predominantly through transcriptional activation of CXCR2 ligands (CXCL1/2/5), establishing this axis as a central mechanism for neutrophil recruitment and subsequent tumor growth in colitis-associated carcinogenesis.

**Figure 6.**
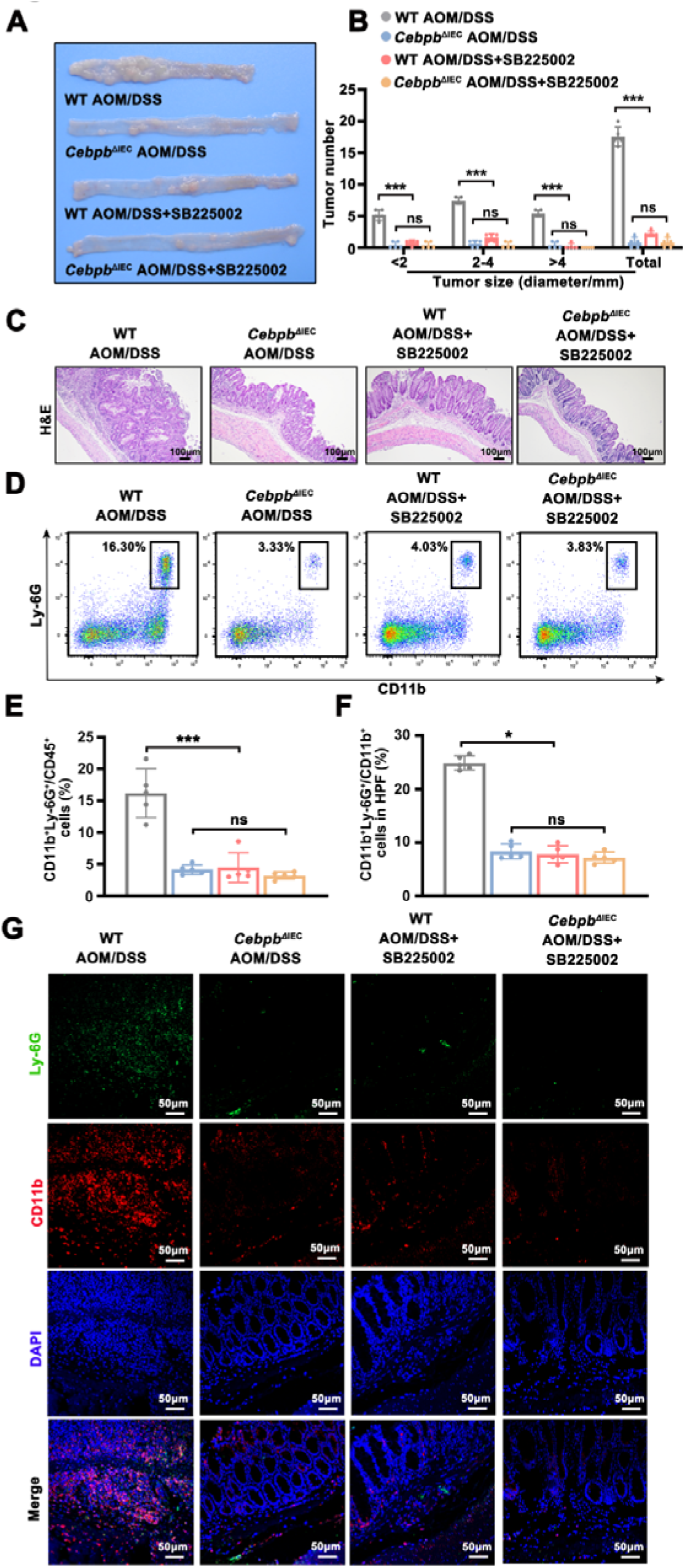
Pharmacological blockade of C/EBPβ-CXCR2 axis reduces neutrophilic inflammation and tumorigenesis in the AOM/DSS CAC model. (A) Colons specimens were harvested from four groups: (i) WT AOM/DSS, (ii) *Cebpb*^ΔIEC^ AOM/DSS, and (iii-iv) SB225002 (CXCR2 antagonist)-treated counterparts (WT or *Cebpb*^ΔIEC^ AOM/DSS mice). (B) Tumor burden quantification (>2 mm by calipers; <2 mm by dissection microscopy; one-way ANOVA with Bonferroni’s correction). n = 5 mice/group. (C) Representative H&E staining in colon tissues from four groups (100×, 100 μm scale). (D-E) Flow cytometry analysis of CD11b^+^Ly-6G^+^ neutrophils among CD45^+^ leukocytes (one-way ANOVA with Bonferroni’s correction). (F-G) Immunofluorescence visualization (200×, 50 μm scale) and ImageJ quantification of Ly-6G^+^ CD11b^+^ cell infiltrate in 5 high-power fields (HPF). n = 5 mice/group. Data are expressed as mean ± SD. ns: not significant, \*\*\**P* < 0.001.

### Immunohistochemical validation and correlation analysis of C/EBP**β**-driven neutrophil infiltration in clinical samples

Immunohistochemical analysis of clinical samples revealed significantly elevated expression of CXCR2 ligands (CXCL1/2/5) and increased neutrophil infiltration (CD66b^+^) in CAC and CRC tissues compared to healthy controls or NAT (Figure 7A-E and Supplementary Figure S7A-E). Strikingly, C/EBPβ levels exhibited a robust positive correlation with CXCR2 ligands, while CXCL1/2/5 expression further correlated with neutrophil infiltration in both CAC and CRC cohorts (Figure 7F-K, and Supplementary Figure S7F-K). These results collectively demonstrate the critical role of the C/EBPβ-CXCL1/2/5 axis in driving neutrophil infiltration during CAC pathogenesis.

**Figure 7.**
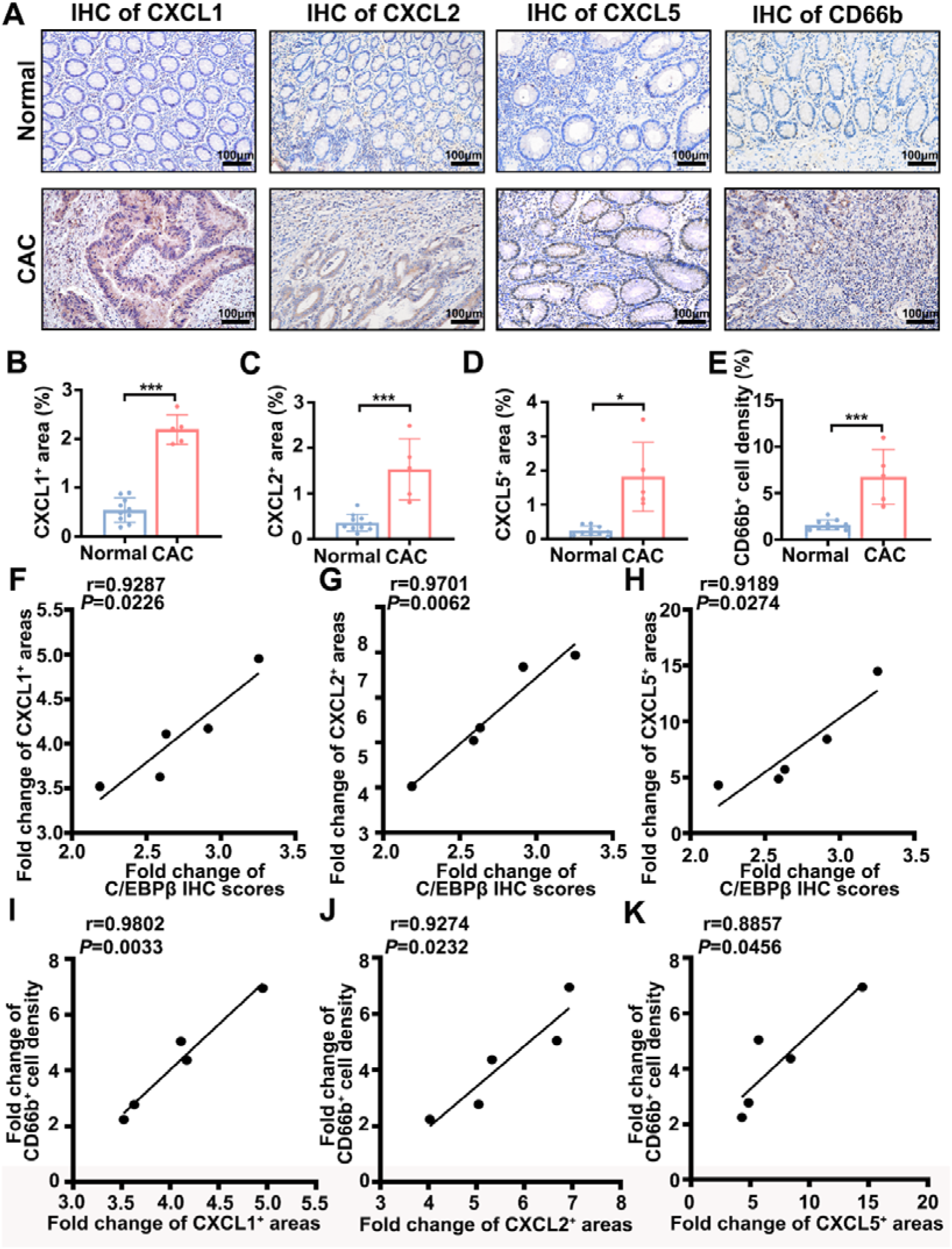
Clinical validation of C/EBPβ-mediated neutrophil recruitment in CAC. (A) Representative IHC staining of CXCL1/2/5, and neutrophil marker CD66b in normal colon (n = 10) vs. CAC (n = 5) (100×, 100 μm scale). (B-E) Quantification of chemokine staining areas and CD66b-positive cell density (CXCL1: unpaired t-test; CXCL2/5/CD66b: Welch’s t-test). (F-K) Pearson correlation analysis: F-H: C/EBPβ IHC scores vs. CXCL1/2/5^+^ areas; I-K: CD66b^+^ cell density vs. CXCL1/2/5^+^ areas. The Pearson correlation coefficient (r) is indicated. Data are expressed as mean ± SD. \**P* < 0.05, \*\*\**P* < 0.001.

## Discussion

Our study establishes intestinal epithelial C/EBPβ as a key transcriptional regulator linking chronic inflammation to colitis-associated carcinogenesis. Through integrated multi-omics analysis, we identified consistent C/EBPβ upregulation across the UC-CAC-CRC continuum, with elevated expression correlating with poorer clinical outcomes. Genetic ablation studies confirmed its functional significance, as epithelial-specific *Cebpb* deletion markedly attenuated both inflammatory responses and tumor development in AOM/DSS models. Mechanistically, we demonstrated that C/EBPβ orchestrates neutrophil recruitment through direct transcriptional activation of CXCR2 ligands (CXCL1/2/5), creating a pro-tumorigenic microenvironment.

Our transcriptomic profiling revealed four additional transcription factors with distinct roles in CAC pathogenesis. FOXQ1 was upregulated in our models, consistent with its known pro-tumorigenic roles in CRC through angiogenic and anti-apoptotic pathways [32–34]. NR3C2 was significantly downregulated, aligning with its tumor suppressive function via AKT/ERK modulation [35, 36] . The specific contributions of FOXQ1 and NR3C2 to CAC require further elucidation. SATB2 depletion, associated with poor prognosis in CRC [37–39], may aggravate CAC through impaired ion homeostasis and microbial dysbiosis [40, 41]. ISX is a novel candidate requiring functional characterization in CAC. These findings position C/EBPβ at the core of a broader transcriptional network governing inflammation-driven carcinogenesis.

C/EBPβ is well-established as a key regulator of immune responses, controlling immune cell development and inflammatory cytokine production [42–44]. Although previous studies have characterized its roles in circulating immune cells during IBD [45, 46], our study reveals its critical epithelial-specific functions in inflammation-driven carcinogenesis. We demonstrate that epithelial C/EBPβ governs neutrophil recruitment to colonic tumors through direct regulation of CXCL1/2/5 expression confirmed by ChIP-qPCR, luciferase assays and EMSA. This epithelial-immune crosstalk is clinically relevant, as evidenced by: (i) correlation with neutrophil accumulation in human CAC/CRC specimens as previous reports [46], and (ii) phenocopy of the *Cebpb*^ΔIEC^ phenotype by CXCR2 inhibition (SB225002).

Notably, the non-additive effect of SB225002 in *Cebpb*^ΔIEC^ mice confirms pathway specificity, consistent with therapeutic outcomes from neutrophil depletion using anti-Gr-1 antibody in the AOM/DSS model [47]. While IL-8 is a well-established transcriptional target of C/EBPβ in humans and binds CXCR2 to drive neutrophil recruitment, mice lack an IL-8 ortholog [48, 49]. Instead, CXCR2 activation in the AOM/DSS model is mediated by functional analogs CXCL1/2/5 [50], establishing a C/EBPβ→CXCL1/2/5→CXCR2→neutrophil recruitment cascade that explains the therapeutic synergy between genetic and pharmacological interventions. Our *in vitro* chemotaxis assays confirmed that SB225002 potently inhibited C/EBPβ-driven neutrophil migration, demonstrating functional conservation of this pathway despite species-specific ligand differences. These findings reveal an epithelial-specific role for C/EBPβ in CAC through CXCR2 ligand-dependent mechanisms, distinct from its immune cell functions [44,45], suggesting tissue-specific therapeutic targeting strategies.

Beyond regulating neutrophil recruitment through the CXCL1/2/5-CXCR2 axis, our study further demonstrates that C/EBPβ coordinates multiple pro-tumorigenic pathways in colitis-associated carcinogenesis. Genetic ablation of C/EBPβ significantly reduced levels of key inflammatory cytokines (IL-1β, IL-6, IL-11, IL-17A, CCL2; Supplementary Figure S4), revealing its broader control of tumor-promoting inflammation. Mechanistically, IL-11 is known to promote the inflammation-cancer transition through STAT3 activation in epithelial/stromal cells [51], IL-17A contributes to colitis-associated tumorigenesis despite its immune origin [52], and CCL2 plays a pivotal role in recruiting tumor-promoting macrophages [53] with its downregulation explaining reduced macrophage infiltration in *Cebpb*^ΔIEC^ mice. Collectively, these findings position C/EBPβ as a master regulator of epithelial-immune crosstalk in CAC. The conserved mechanism across murine models and human disease, and druggable nature of multiple downstream effectors (CXCR2 ligands, IL-11/STAT3, CCL2) highlight promising therapeutic opportunities, particularly through combinatorial targeting strategies or direct epithelial C/EBPβ inhibition for CAC treatment. Our work provides both mechanistic insights for understanding inflammation-driven tumorigenesis and actionable targets for CAC intervention.

## Supplementary Information

Supplementary Data, Figures, and Tables are available at Supplementary information.

## Ethics declarations

All human participant research was conducted in accordance with the Declaration of Helsinki and approved by the Institutional Review Board of Jinling Hospital (No. 2021DZSKT-YBB-008). Written informed consent was obtained from all participants prior to study enrollment. Animal experiments were conducted in compliance with ARRIVE guidelines and approved by the Institutional Animal Care and Use Committee of Nanjing University (IACUC-2006015).

## Availability of Data and Materials

All data are available in the article and its supplementary materials. Raw transcriptome sequencing data have been deposited in the NCBI Gene Expression Omnibus (GEO) database under accession series GSE264497.

## Author Contributions

J. Chen and Z. Huang conceived and supervised the study. Mingyue Li, Xintong Wang, Wenjie Hu, Xiaohui Cheng, Qi sun performed experiments. Xintong Wang and Mingyue Li conducted data analysis. Yongjie Wu, and Xintong Wang provided analytical tools. J. Chen led manuscript drafting, rigorous revisions, and final integration. Mingyue Li, and Xintong Wang assisted in preliminary drafting. J. Chen and Z. Huang performed critical revisions and finalized the manuscript. All authors approved the final version.

## Acknowledgments

This work was supported by the National Natural Science Foundation of China (Grant numbers: 31771550, 31870821 and 81972267) and Jiangsu Provincial Science and Technology Plan Special Fund (BZ2024052 and BM2023008).

