## Supplementary material for "Intestinal Epithelial C/EBPβ Deficiency Impairs Colitis-Associated Tumorigenesis by Disrupting CXCL1/CXCL2/CXCL5-CXCR2-Mediated Neutrophil Infiltration": Supplementary information.docx

**Supplementary Figures**

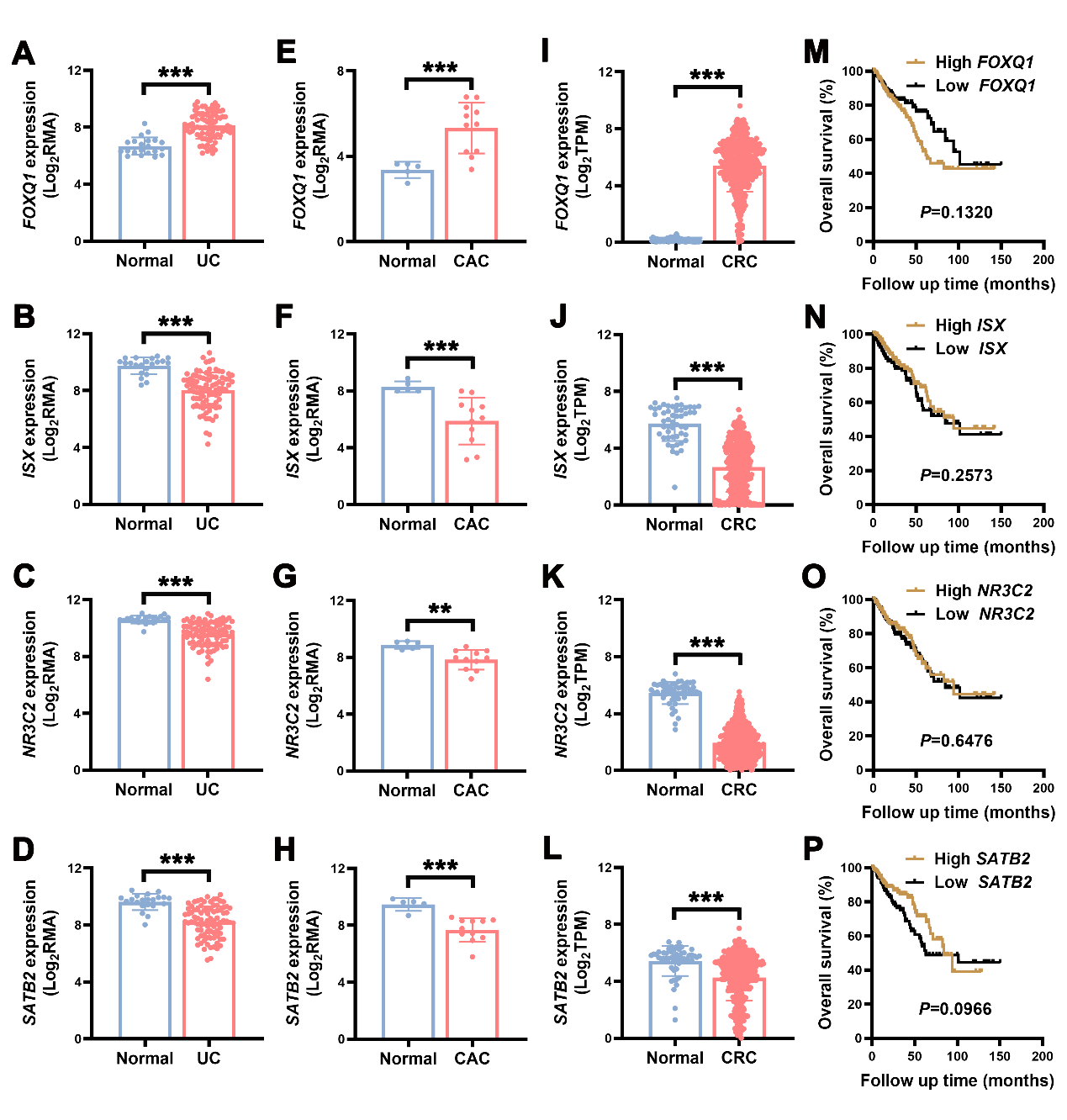

**Figure S1. Gene expression of *FOXQ1*, *ISX*, *NR3C2*, and *SATB2* in UC, CAC and CRC, and their prognostic correlation in CRC.** (A-D) mRNA levels of *FOXQ1*, *ISX*, *NR3C2*, and *SATB2* were examined in normal (n = 21) and UC patients (n = 87), utilizing the GSE87466 dataset (Welch’s t-test (two-tailed) in panel A or two-tailed Mann-Whitney U test in panels B-D). (E-H) mRNA levels of *FOXQ1*, *ISX*, *NR3C2*, and *SATB2* were investigated in normal (n = 5) and CAC patients (n = 11) from the GSE37283 dataset (Welch’s t-test (two-tailed) in panels E-F or Unpaired two-tailed Student’s t-test in panels G-H). (I-L) mRNA levels of *FOXQ1*, *ISX*, *NR3C2*, and *SATB2* were assessed in normal (n = 51) and CRC patients (n = 383) using the TCGA-COAD/READ dataset (Mann-Whitney test). (M-P) Kaplan-Meier survival curves for 376 CRC patients, stratified by mRNA expression levels of *FOXQ1*, *ISX*, *NR3C2*, and *SATB2*, were generated from the TCGA-COAD/READ dataset. Data are expressed as mean ± SD. ***P* < 0.01, ****P* < 0.001.

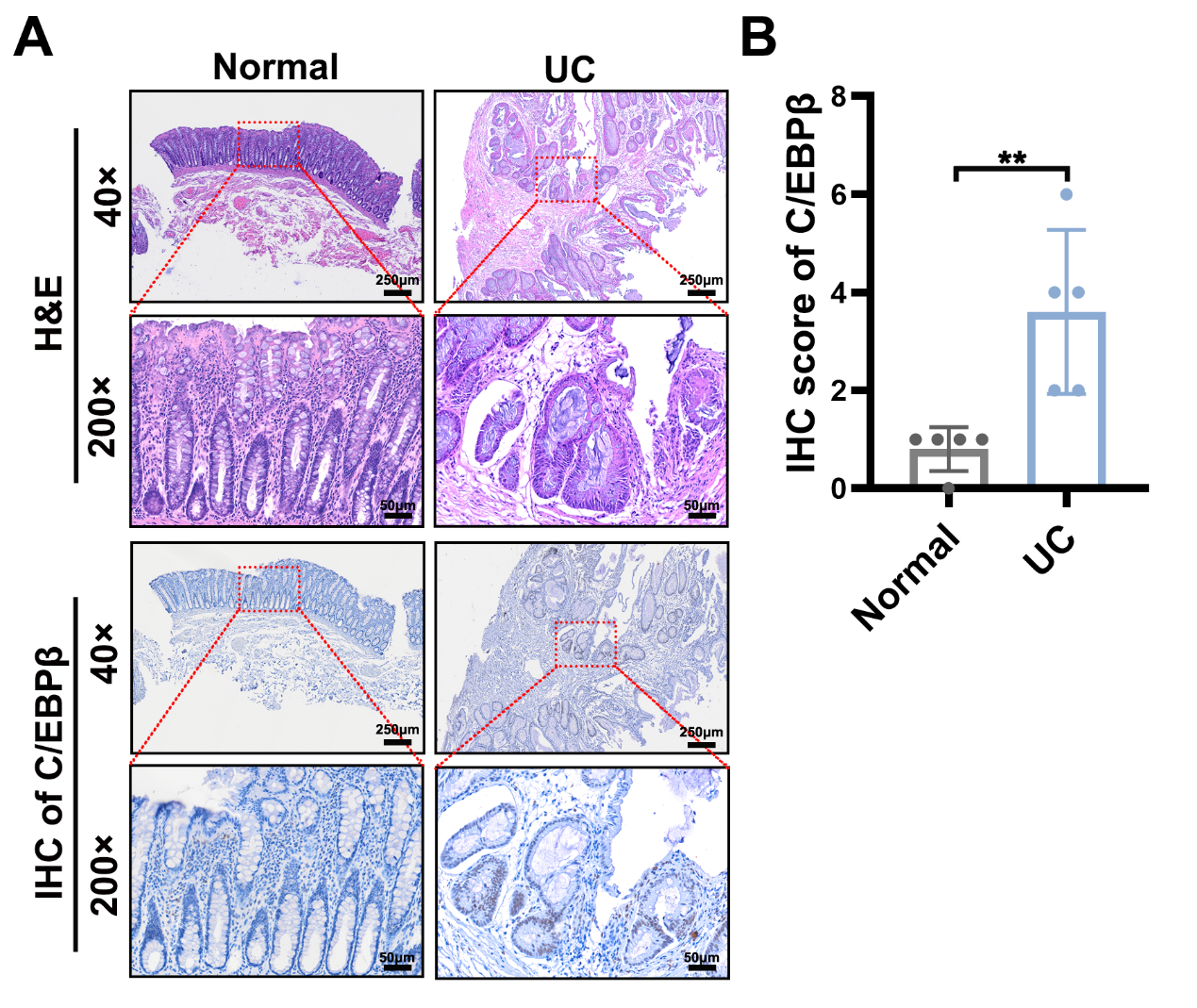

**Figure S2.** **Clinical significance of elevated C/EBPβ in ulcerative colitis (UC).** (A) Representative hematoxylin and eosin (H&E) staining and C/EBPβ immunohistochemistry (IHC) in normal colon and UC colon specimens. Scale bar: 250 μm (40× magnification). (B) Quantitative analysis of C/EBPβ IHC staining intensity (Normal: n = 5; UC: n = 5). Data are expressed as mean ± SD. (***P* < 0.01 by unpaired two-tailed Student's t-test).

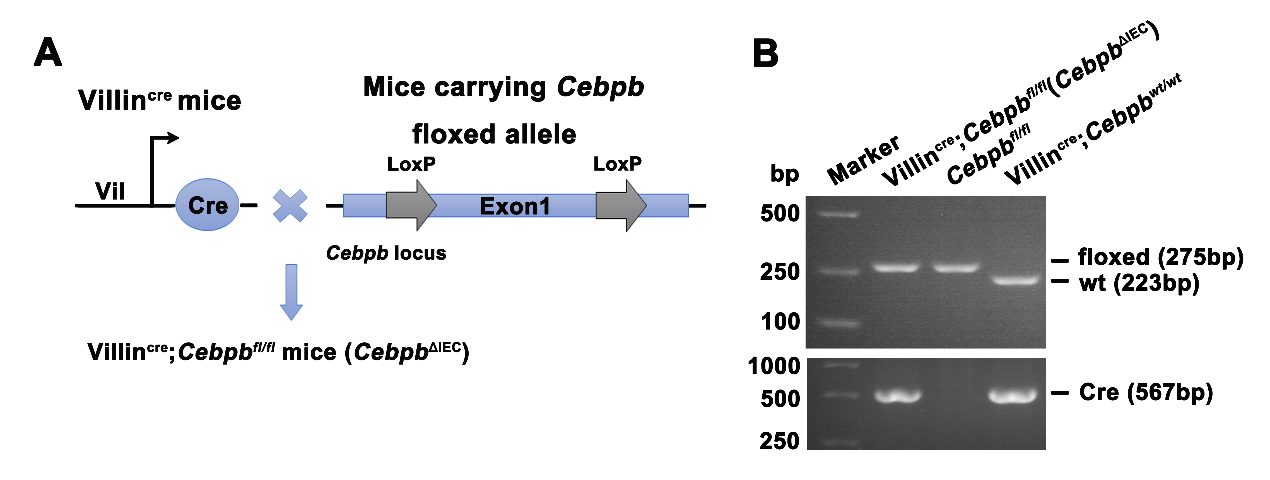

**Figure S3. Generation of intestinal epithelial cell-specific *Cebpb* knockout mice.** (A) A schematic illustration shows the generation of *Cebpb*^ΔIEC^ mice through cross-breeding *Villin*^Cre^ mice with *Cebpb*^fl/fl^ mice. (B) Genotypic confirmation was performed using PCR on tail-tip DNA and agarose gel electrophoresis, assessing both the floxed allele (Top) and the Cre-mediated recombined allele (Bottom). Representative images from three independent experiments are presented.

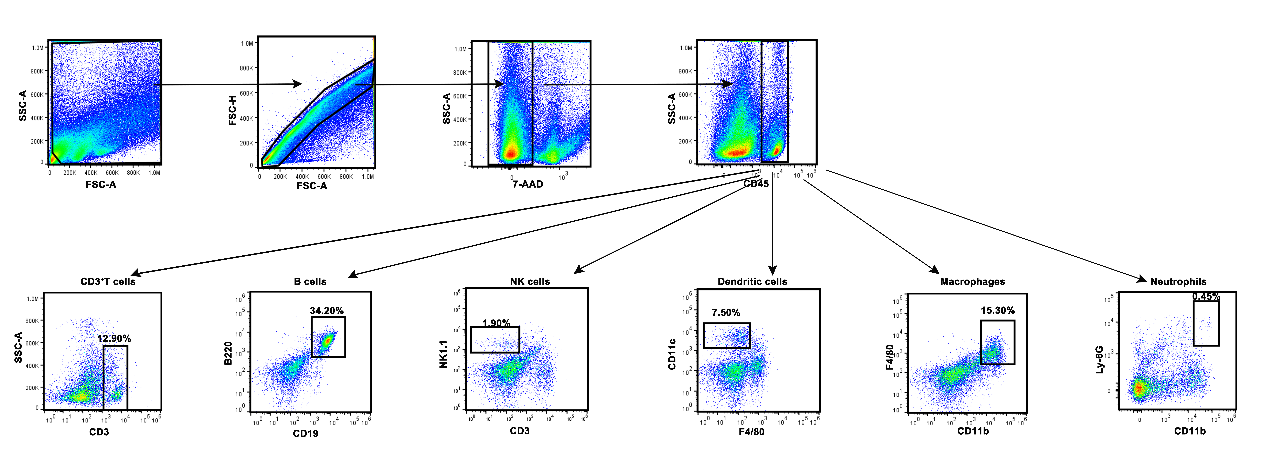

**Figure S4. Flow cytometry gating strategy for colonic immune cells in mice.** The gating strategy encompasses the identification of CD3^+^ T cells, CD19^+^B220^+^ B cells, CD3^-^NK1.1^+^ NK cells, F4/80^-^CD11c^+^ dendritic cells, CD11b^+^F4/80^+^ macrophages and CD11b^+^Ly-6G^+^ neutrophils within the CD45^+^ immune cell population of the colon in mice.

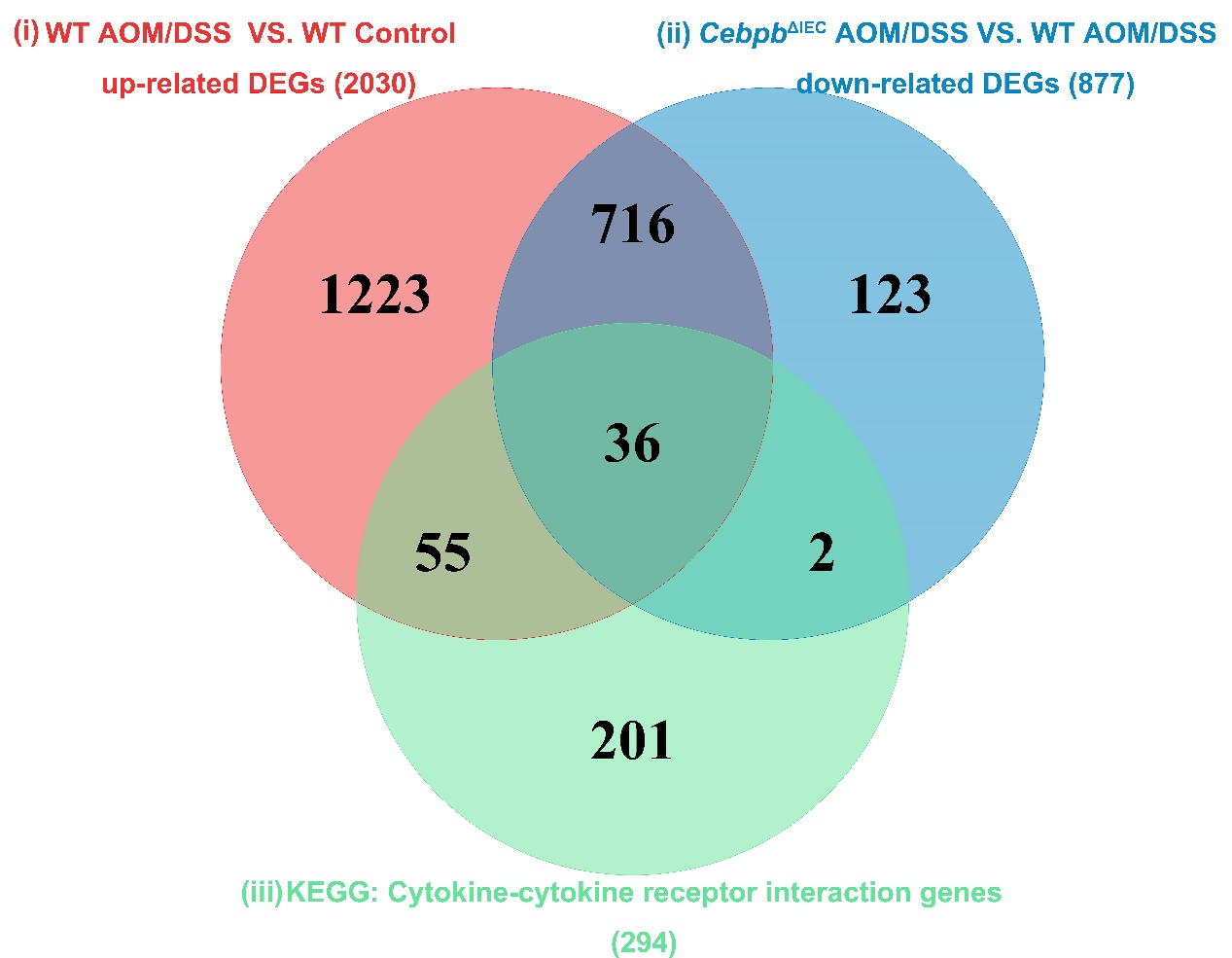

**Figure S5. Venn diagram showing intersecting gene sets:** (i) upregulated in WT AOM/DSS vs controls (pink), (ii) downregulated in *Cebpb*^ΔIEC^ vs WT AOM/DSS (blue), and (iii) cytokine-cytokine receptor pathway genes from GSEA (green).

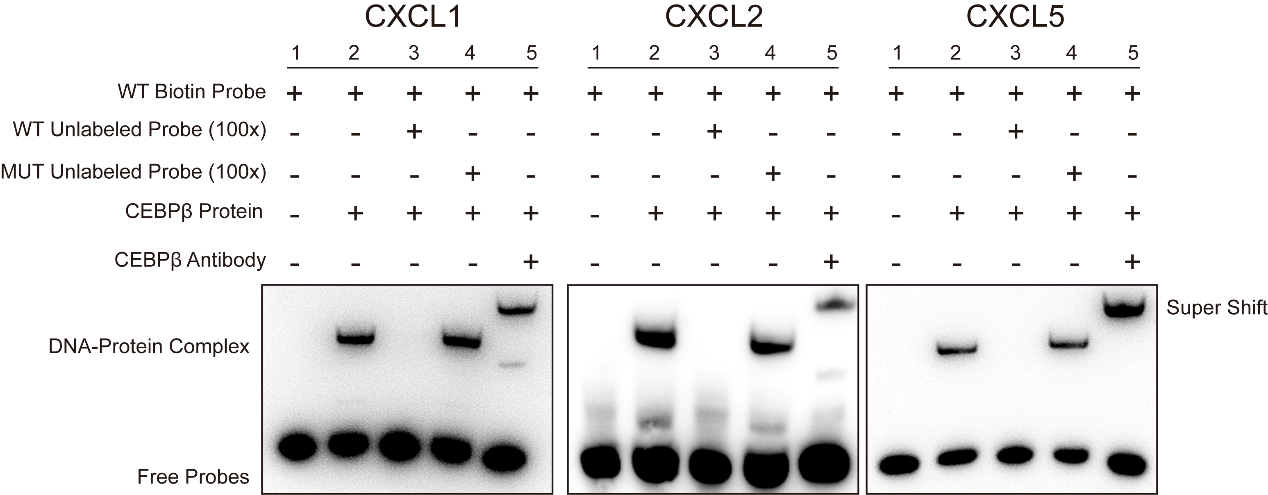

**Figure S6. Electrophoretic mobility shift assay (EMSA) analysis of CEBPβbinding to CXCL1, CXCL2, and CXCL5 promoter regions.** All reactions contained biotin-labeled probes, with lane 1 as probe-only control and lane 2 showing C/EBPβ-probe complexes. Specific binding was confirmed by competition assays using 100-fold molar excess of unlabeled wild-type probes (lane 3) versus mutated non-specific competitors (MUT, lane 4). Supershift assays (lane 5) with C/EBPβ-specific antibody further verified binding specificity. Mutated probes (MUT) failed to compete for C/EBPβ binding, demonstrating sequence specificity**.**

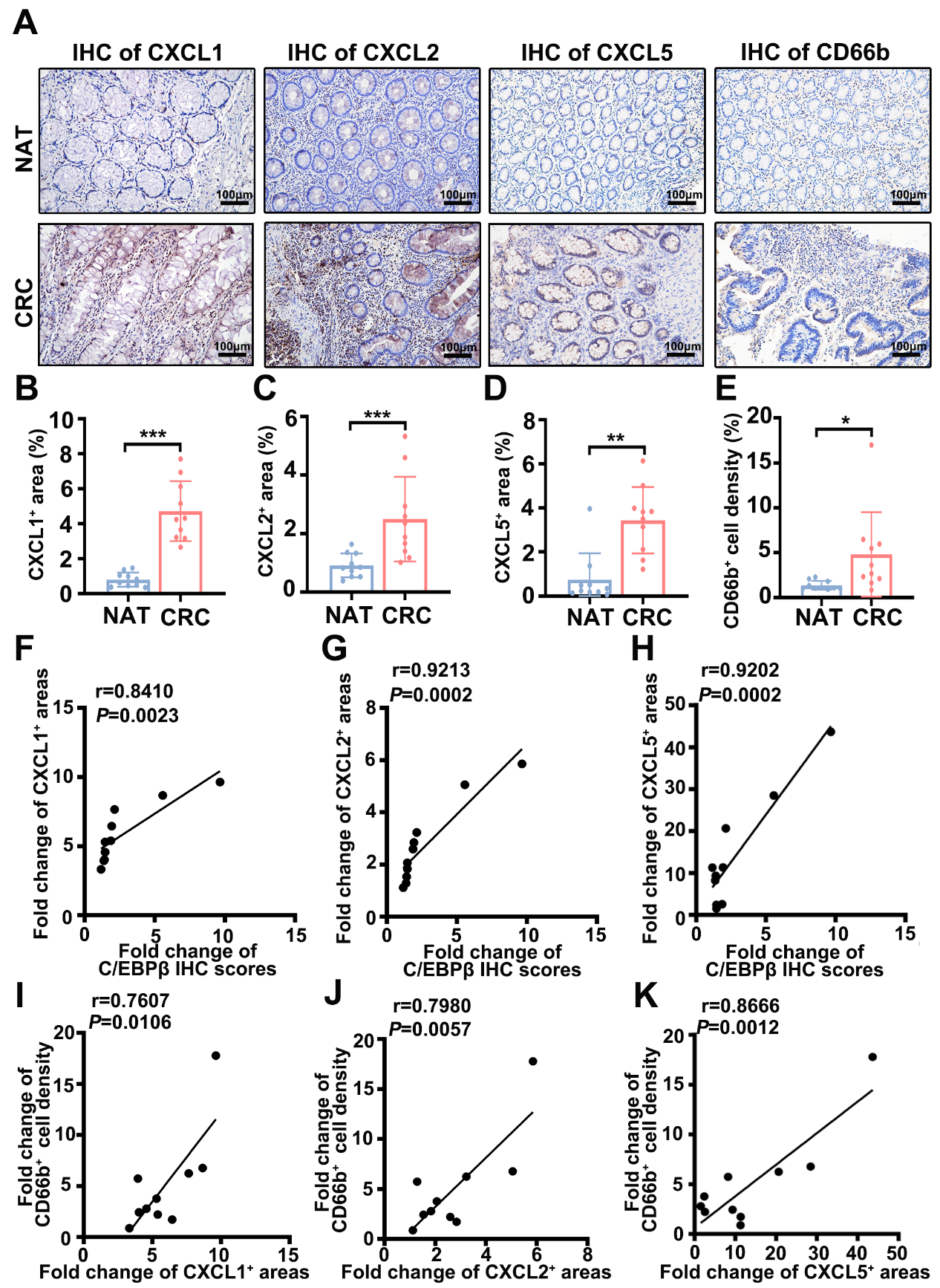

**Figure S7. Clinical validation of C/EBPβ-mediated chemotaxis in CRC samples.** (A) IHC staining of CXCL1, CXCL2, CXCL5, and CD66b in CRC tissues and NAT (n = 10) with a magnification of 100× and a scale bar of 100 μm. (B-E) Quantification of positive areas for CXCL1 (B), CXCL2 (C), CXCL5 (D), and CD66b (E) in NAT and CRC samples using the Welch’s t-test (two-tailed) in panels B, C, and E or two-tailed Mann-Whitney U test in panel D. (F-H) Pearson’s correlation scatter plots elucidate the relationship between the fold changes in C/EBPβ IHC scores and the positive areas for CXCL1, CXCL2, and CXCL5 in NAT and CRC tissues. (I-K) Pearson’s correlation scatter plots depict the relationship between the fold changes of the positive density for CD66b cell and the positive areas for CXCL1, CXCL2, CXCL5, and CD66b in NAT and CRC tissues. The Pearson correlation coefficient (r) is indicated. n = 10 patients. Data are expressed as mean ± SD. ***P* < 0.01, ****P* < 0.001.

**Supplementary Tables**

**Supplementary Table S1. Clinicopathological characteristics of 87 UC patients from GEO dataset GSE87466 used for bioinformatics analysis.**

| **Variables** | **Number (n)** | **Proportion (%)** |
| --- | --- | --- |
| ***All cases*** | 87 | 100 |
| ***Age(yr)*** |  |  |
| ≤ 60 | 75 | 86.2 |
| > 60 | 12 | 13.8 |
| ***Disease extent*** |  |  |
| Limited | 60 | 69.0 |
| Extensive | 27 | 31.0 |

**Supplementary Table S2. Clinicopathological characteristics of 11 CAC patients and 5 healthy controls from GEO dataset GSE37283 used for bioinformatics analysis.**

| **Variables** | **CAC patients** | **Healthy individuals** |
| --- | --- | --- |
| ***All cases*** | 11 | 5 |
| ***Gender*** |  |  |
| Male | 11 | 5 |
| Female | 0 | 0 |
| ***Age (Mean)*** | 46.9 | 46 |
| ***Dysplasia histology*** |  | NA |
| Low grade dysplasia | 4 |  |
| High grade dysplasia | 3 |  |
| Adenocarcinoma | 4 |  |

**Supplementary Table S3. Clinicopathological characteristics of 383 CRC patients in the TCGA-COAD/READ cohort analyzed through bioinformatics approaches.**

|  | **Tumor tissues from**  **CRC patients** | **Normal adjacent tissues from CRC patients** |
| --- | --- | --- |
| ***All cases*** | 383 | 51 |
| ***Gender*** |  |  |
| Male | 208 | 23 |
| Female | 171 | 28 |
| Unknown | 4 | 0 |
| ***Age(yr)*** |  |  |
| ≤ 60 | 144 | 12 |
| > 60 | 233 | 39 |
| Unknown | 6 | 0 |
| ***Stage (TNM)*** |  |  |
| T1 | 57 | 2 |
| T2 | 137 | 7 |
| T3 | 114 | 36 |
| T4 | 52 | 6 |
| Unknown | 23 | 0 |
| ***Lymphatic metastasis*** |  |  |
| Yes | 170 | 17 |
| No | 207 | 34 |
| Unknown | 6 | 0 |
| ***Distant metastasis*** |  |  |
| Yes | 117 | 9 |
| No | 255 | 35 |
| Unknown | 11 | 7 |

**Supplementary Table S4. Tissue microarray sample information from 180 CRC cases (H&E and C/EBPβ IHC analysis).**

|  | **Number(n)** | **Proportion (%)** |
| --- | --- | --- |
| ***All cases*** | 180 | 100 |
| ***Gender*** |  |  |
| Male | 92 | 51.1 |
| Female | 88 | 48.9 |
| ***Age(yr)*** |  |  |
| ≤ 60 | 37 | 20.6 |
| > 60 | 143 | 79.4 |
| ***Stage (TNM)*** |  |  |
| T1 | 14 | 7.8 |
| T2 | 99 | 55.0 |
| T3 | 64 | 35.5 |
| T4 | 3 | 1.7 |
| ***Histological type*** |  |  |
| Well differentiated | 28 | 15.6 |
| Moderately differentiated | 107 | 59.4 |
| Poorly differentiated | 45 | 25.0 |
| ***Tumor size*** |  |  |
| ≤5 cm | 104 | 59.3 |
| >5 cm | 76 | 40.7 |
| ***Lymphatic metastasis*** |  |  |
| Yes | 67 | 46.3 |
| No | 113 | 53.7 |

**Supplementary Table S5. Clinical data of UC (n = 5), CAC cohort (n = 5), and healthy controls (n = 10) analyzed by H&E staining, IHC (C/EBPβ, CXCL1-5, CD66b), and C/EBPβ qRT-PCR.**

| **Variables** | **UC patients** | **CAC patients** | **Healthy individuals** |
| --- | --- | --- | --- |
| ***Number of cases*** | 5 | 5 | 10 |
| ***Age*** (≤ 45 / > 45) | 1/4 | 3/2 | 5/5 |
| ***Gender*** (male / female) | 3/2 | 2/3 | 6/4 |
| ***Histological type*** (Well differentiated/ Poorly differentiated) | NA | 4/1 | NA |

**Supplementary Table S6. Clinical data of 64 CRC cases analyzed for C/EBPβ expression (Western blot/qRT-PCR) and protein localization (IHC: C/EBPβ, CXCL1/2/5, CD66b).**

|  | **Number (n)** | **Proportion (%)** |
| --- | --- | --- |
| ***All cases*** | 64 | 100 |
| ***Gender*** |  |  |
| Male | 38 | 59.4 |
| Female | 26 | 40.6 |
| ***Age(yr)*** |  |  |
| ≤ 60 | 27 | 42.2 |
| > 60 | 37 | 57.8 |
| ***Stage (TNM)*** |  |  |
| T1 | 0 | 0 |
| T2 | 13 | 20.3 |
| T3 | 30 | 46.9 |
| T4 | 21 | 32.8 |
| ***Histological type*** |  |  |
| Well differentiated | 9 | 14.1 |
| Moderately differentiated | 47 | 73.4 |
| Poorly differentiated | 8 | 12.5 |
| ***Lymphatic metastasis*** |  |  |
| Yes | 28 | 43.8 |
| No | 36 | 56.2 |

**Supplementary Table S7. siRNA sequences and EMSA probes used in this study.**

| **Name** | | **Sequence (5’-3’)** |
| --- | --- | --- |
|  | Homo-*CEBPB* siRNA sense | GCCUGCCUUUAAAUCCAUGTT |
|  | Homo-*CEBPB* siRNA antisense | CAUGGAUUUAAAGGCAGGCTT |
|  | Negative control siRNA for homo-*CEBPB* sense | UUCUCCGAACGUGUCACGUTT |
|  | Negative control siRNA for homo-*CEBPB* antisense | ACGUGACACGUUCGGAGAATT |
|  | *CXCL1* WT biotin probe sense | Biotin-TGGAACTTTCAAAATCCCCT |
|  | *CXCL1* WT biotin probe antisense | Biotin-AGGGGATTTTGAAAGTTCCA |
|  | *CXCL1* WT unlabeled probe sense | TGGAACTTTCAAAATCCCCT |
|  | *CXCL1* WT unlabeled probe antisense | AGGGGATTTTGAAAGTTCCA |
|  | *CXCL1* MUT unlabeled probe sense | TGGAAACGGTCCCCACCCCT |
|  | *CXCL1* MUT unlabeled probe antisense | AGGGGTGGGGACCGTTTCCA |
|  | *CXCL2* WT biotin probe sense | Biotin- TAGAAGTGGTGTTTCACAACCTTACTGATA |
|  | *CXCL2* WT biotin probe antisense | Biotin- TATCAGTAAGGTTGTGAAACACCACTTCTA |
|  | *CXCL2* WT unlabeled probe sense | TAGAAGTGGTGTTTCACAACCTTACTGATA |
|  | *CXCL2* WT unlabeled probe antisense | TATCAGTAAGGTTGTGAAACACCACTTCTA |
|  | *CXCL2* MUT unlabeled probe sense | TAGAAGTGGTAGGGTCTCCTCTTACTGATA |
|  | *CXCL2* MUT unlabeled probe antisense | TATCAGTAAGAGGAGACCCTACCACTTCTA |
|  | *CXCL5* WT biotin probe sense | Biotin-CACTGTCTTGCTCCATGGCAG |
|  | *CXCL5* WT biotin probe antisense | Biotin-CTGCCATGGAGCAAGACAGTG |
|  | *CXCL5* WT unlabeled probe sense | CACTGTCTTGCTCCATGGCAG |
|  | *CXCL5* WT unlabeled probe antisense | CTGCCATGGAGCAAGACAGTG |
|  | *CXCL5* MUT unlabeled probe sense | CACTGTCAATTTTTATGGCAG |
|  | *CXCL5* MUT unlabeled probe antisense | CTGCCATAAAAATTGACAGTG |

**Supplementary Table S8. Antibodies for flow cytometry, western blot, immunofluorescence and immunohistochemistry.**

| **Antibody name** | **Corporation name** | **Cat. NO.** | **Dilution ratio (WB)** | **Dilution ratio**  **(IHC/IF)** | **Dilution ratio**  **(FC)** |
| --- | --- | --- | --- | --- | --- |
| C/EBPβ | Santa cruz | sc-7962 | / | 1:1000 | / |
| C/EBPβ | Abcam | Ab32358 | 1:2000 | / | / |
| CXCL1 | Proteintech | 12335-1-AP | / | 1:200 | / |
| CXCL2 | Abbexa | Abx3778774 | / | 1:200 | / |
| CXCL5 | Affinity Biosciences | DF9919 | / | 1:500 | / |
| CD66b | Abcam | Ab214175 | / | 1:200 | / |
| CD11b | Abcam | Ab133357 | / | 1:2000 | / |
| Ly-6G | Biolegend | 127601 | / | 1:100 | / |
| GAPDH | Proteintech | HRP-60004 | 1:10000 | / | / |
| CD45 | Abcam | Ab40763 | 1:2000 | 1:200 | / |
| Pan Cytokeratin | Abcam | ab7753 | 1:2000 | / | / |
| α-SMA | Boster | BM0002 | 1:2000 | / | / |
| Goat anti-mouse IgG-HRP | Jackson ImmunoResearch | 115-035-003 | 1:2000 | / | / |
| Goat anti-rabbit IgG-HRP | Jackson ImmunoResearch | 111-035-003 | 1:2000 | / | / |
| Alexa Fluor 488-conjugated donkey anti-rat | Invitrogen | A21208 | / | 1:200 | / |
| Alexa Fluor 546-conjugated donkey anti-rabbit | Invitrogen | A10040 | / | 1:200 | / |
| 7-AAD Viability Staining Solution | Biolegend | 420404 | / | / | 1:100 |
| FITC anti-mouse CD45 | Biolegend | 103108 | / | / | 1:200 |
| APC anti-mouse CD3 | Biolegend | 100236 | / | / | 1:20 |
| PE anti-mouse CD19 | Biolegend | 115508 | / | / | 1:80 |
| PE/Cyanine7 anti-mouse NK-1.1 | Biolegend | 156514 | / | / | 1:20 |
| Brilliant Violet 421™ anti-mouse/human CD45R/B220 | Biolegend | 103251 | / | / | 1:20 |
| PE/Cyanine7 anti-mouse CD11b | Biolegend | 101216 | / | / | 1:80 |
| APC anti-mouse CD11c | Biolegend | 117310 | / | / | 1:80 |
| PE anti-mouse F4/80 | Biolegend | 123110 | / | / | 1:20 |
| Brilliant Violet 421™ anti-mouse Ly-6G | Biolegend | 108445 | / | / | 1:20 |

**Supplementary Table S9. Primer sequences used in this study.**

| **Name** | | **Primer Sequence (5’-3’)** |
| --- | --- | --- |
| Homo*-CXCL1* Forward (ChIP-qPCR) | | AGTGACAACCAGTGCCGTAT |
| Homo*-CXCL1* Reverse (ChIP-qPCR) | | GGTGGCAACAGATTA |
| Homo*-CXCL2* Forward (ChIP-qPCR) | | GCCATACCTTCATAACC |
| Homo*-CXCL2* Reverse (ChIP-qPCR) | | TAATAGTGCCTGCCTCA |
| Homo*-CXCL5* Forward (ChIP-qPCR) | | GAAAGCCTATGTGTCATTCCAATAC |
|  | Homo*-CXCL5* Reverse (ChIP-qPCR) | GCACTGAAGGTTTAAGGGTTTTC |
|  | Homo*-ACTB* Forward | GACTGCTGTCACCTTCACCGTTC |
|  | Homo*-ACTB* Reverse | GACTTAGTTGCGTTACACCCTTTCTTG |
|  | Mus-*Actb* Forward | GTGAAAAGATGACCCAGATCAT |
|  | Mus-*Actb* Reverse | GCTTCTCTTTGATGTCACGCACGAT |
|  | Homo*-CEBPB* Forward | AAGCACAGCGACGAGTACAA |
|  | Homo*-CEBPB* Reverse | ACAGCTGCTCCACCTTCTTC |
|  | Mus-*Cebpb* Forward | ACTTCCTCTCCGACCTCTTC |
|  | Mus-*Cebpb* Reverse | GCTCACGTAACCGTAGTCG |
|  | Homo*-CXCL1* Forward | AAGAACATCCAAAGTGTGAACG |
|  | Homo*-CXCL1* Reverse | CACTGTTCAGCATCTTTTCGAT |
|  | Mus-*Cxcl1* Forward | GGCTGGGATTCACCTCAAGAACATC |
|  | Mus-*Cxcl1* Reverse | TGAGTGTGGCTATGACTTCGGTTTG |
|  | Homo*-CXCL2* Forward | AAGTGTGAAGGTGAAGTCCC |
|  | Homo*-CXCL2* Reverse | TCTGCCCATTCTTGAGTGTG |
|  | Mus-*Cxcl2* Forward | TCAATGCCTGAAGACCCTG |
|  | Mus-*Cxcl2* Reverse | CCTTGAGAGTGGCTATGACTTC |
|  | Homo*-CXCL5* Forward | TCTGCAAGTGTTCGCCATAG |
|  | Homo*-CXCL5* Reverse | TTCCACCGTCCAAAATTTTCTG |
|  | Mus-*Cxcl5* Forward | TGCCCCTTCCTCAGTCATAG |
|  | Mus-*Cxcl5* Reverse | AGGGATCACCTCCAAATTAGC |
|  | Mus-Cxcr2 Forward | ATGCCCTCTATTCTGCCAGAT |
|  | Mus-Cxcr2 Reverse | GTGCTCCGGTTGTATAAGATGA |

**Supplementary Table S10. Histopathological classification of colorectal tumors in AOM/DSS-treated WT versus *Cebpb*^ΔIEC^ mice (n = 5/group).**

| **Group** | **Low-grade adenoma** | **High-grade adenoma** | **Invasive carcinomas** |
| --- | --- | --- | --- |
| **WT+ AOM/DSS** | 0 | 80% | 20% |
| ***Cebpb*^ΔIEC^ +AOM/DSS** | 20% | 0 | 0 |

**Supplementary Table S11. 36 consensus DEGs from WT AOM/DSS-upregulated, *Cebpb*^ΔIEC^-downregulated, and GSEA** **cytokine-cytokine receptor pathway analyses.**

| **Serial number** | **Gene name** |
| --- | --- |
| 1 | *Il17a* |
| 2 | *Csf3* |
| 3 | *Cxcl5* |
| 4 | *Il11* |
| 5 | *Il17f* |
| 6 | *Cxcr2* |
| 7 | *Il6* |
| 8 | *Cxcl1* |
| 9 | *Il1a* |
| 10 | *Ccl7* |
| 11 | *Gdf11* |
| 12 | *Tnfrsf11b* |
| 13 | *Cxcl2* |
| 14 | *Ccl4* |
| 15 | *Tnfsf18* |
| 16 | *Inhba* |
| 17 | *Tnfrsf19* |
| 18 | *Il1b* |
| 19 | *Tnfrsf9* |
| 20 | *Il33* |
| 21 | *Cxcl9* |
| 22 | *Ccl3* |
| 23 | *Ccl12* |
| 24 | *Bmp7* |
| 25 | *Osm* |
| 26 | *Il20ra* |
| 27 | *Ccr8* |
| 28 | *Il23a* |
| 29 | *Ccl2* |
| 30 | *Eda2r* |
| 31 | *Csf3r* |
| 32 | *Cxcl14* |
| 33 | *Ccr5* |
| 34 | *Il1r2* |
| 35 | *Il1rl1* |
| 36 | *Tnfsf4* |

**Supplementary Table S12. Histopathological classification of colorectal tumors in SB225002-treated AOM/DSS models: *Cebpb*^ΔIEC^ versus wild-type (n = 5/group).**

| **Group** | **Low-grade adenoma** | **High-grade adenoma** | **Invasive carcinomas** |
| --- | --- | --- | --- |
| **WT+AOM/DSS+SB225002** | 40% | 0 | 0 |
| ***Cebpb*^ΔIEC^ +AOM/DSS+SB225002** | 20% | 0 | 0 |
